# Memory network activity flow failures in temporal and frontal lobe epilepsy

**DOI:** 10.1101/2025.11.27.690997

**Authors:** Donna Gift Cabalo, Jordan DeKraker, Ke Xie, Thaera Arafat, Jessica Royer, Raúl Rodríguez-Cruces, Alexander Ngo, Ella Sahlas, Raluca Pana, Elizabeth Jefferies, Alexander James Barnett, Lorenzo Caciagli, Samantha Audrain, R. Nathan Spreng, Andrea Bernasconi, Neda Bernasconi, Boris C. Bernhardt

## Abstract

Declarative memory emerges from distributed brain networks, yet how large-scale network organization constrains memory function remains unclear. Intrinsic connectivity mirrors task activity, suggesting intrinsic architecture governs memory states. Temporal and frontal lobe epilepsy (TLE, FLE), differentially affecting episodic and semantic systems, provide complementary models of network disruption. Here, we combine resting-state and task fMRI in 87 healthy controls, 33 TLE and 20 FLE patients using activity flow mapping, a generative framework predicting task activation from intrinsic connectivity. Activity flow mapping reconstructs memory activations in controls but shows reduced prediction accuracy in patients, co-occurring with abnormal task activations, intrinsic dysconnectivity, and episodic impairment without significant semantic impairment. Activity flow mapping accuracy, but not activation or connectivity alone, captures clinical variation and is associated with intrinsic functional and microstructural network features. Activity flow mapping connects intrinsic brain architecture to declarative memory and identifies altered network communication as a correlate of memory dysfunction and a potential intervention target.

## Introduction

Human memory emerges from interactions within distributed, modular, and hierarchical brain networks that integrate specialized regional computations into coherent cognitive states, rather than from isolated cortical loci. ^1–3^ Although episodic and semantic memory, both forms of declarative memory, rely on common substrates in the medial and lateral temporal lobes as well as posterior and frontal regions, ^4–11^ there is also evidence for distinct pathways. Indeed, it is well established that the hippocampus and medial temporal lobe (MTL) in particular contribute to episodic memory function ^12–14^ while semantic memory more critically involves more integrative anterior temporal lobe (ATL) *hub* and modality-specific *spoke* regions. ^15–17^ Concomitant convergence and divergence of these systems ^18^ facilitate flexible memory processing, supporting recall of specific episodic experience and general semantic knowledge.

Previous work suggests that intrinsic architecture constrains cognitive states, with resting-state functional connectivity (rsFC) closely mirroring task-evoked activation and providing a scaffold for task-related brain activity. ^19,20^ Importantly, although brain activity reorganizes across cognitive contexts, these state-dependent patterns remain constrained by stable, person-specific intrinsic connectivity, enabling the prediction of task-evoked responses from resting-state network patterns. Consistent with this view, computational and empirical work indicates that task-evoked activity can be understood to reflect the propagation of activation over intrinsic functional pathways, linking distributed connectivity to localized task responses and cognitive performance. ^20–22^ Activity flow mapping (AFM) ^21,23^ formalizes this principle by leveraging rsFC and task activations to model how distributed brain networks support cognition. Specifically, AFM estimates task-evoked activation in a target region based on activity from other regions, weighted by their rsFC. ^21,23^ AFM uses interpretable correlational estimates of connectivity and generalized linear models (GLM) of task activity, incorporating confound control and empirical validation to test predictions against fMRI data. AFM has demonstrated utility in healthy aging, ^21^ neurodegenerative conditions, ^24^ and schizophrenia.^25^ In these conditions, altered rsFC was shown to predict both dysfunctional activations and brain-behaviour relationships. ^25^ Computational simulations further revealed that targeted modifications to connectivity could normalize dysfunctional activity patterns. ^25^ Taken together, these findings indicate that AFM framework offers a powerful approach for understanding how intrinsic connectivity organizes memory-related activity in the healthy brain, providing a normative reference against which disease-related alterations can be interpreted.

Temporal lobe epilepsy (TLE) and frontal lobe epilepsy (FLE), two of the most common pharmaco-resistant epilepsies, provide complementary and compelling human disease models for examining how disruption of temporal and frontal systems perturbs distributed episodic and semantic memory circuits. ^26^ Declarative memory deficits are key features in both TLE and FLE, ^27–33^ and studying these deficits offers a systems-level framework to conceptualize common and distinct pathways implicated in episodic and semantic memory processing. TLE is typically associated with episodic memory deficits, ^27,28^ which are thought to be related to structural compromise in the MTL, ^34–36^ as well as in more distant temporal and extratemporal regions. ^37,38^ While sometimes observed in TLE patients, semantic memory losses appear milder and overall, more variable. ^29,30^ Similarly, patients undergoing selective MTL resections often exhibit substantial episodic memory impairments, ^39,40^ whereas semantic memory performance tends to be preserved. ^41,42^ This apparent resilience may reflect the broader and more distributed neural architecture supporting semantic memory, as opposed to the more MTL-dependent systems implicated in episodic memory. ^43,44^ However, contradictory evidence points to measurable semantic impairments in some TLE patients who present with structural compromise in lateral and ATL regions, ^45,46^ including those who have undergone unilateral ATL resections to treat their seizures. ^46,47^

Contrary to TLE, FLE is characterized by seizures originating often from a broad spectrum of regions across the frontal lobe,^48,49^ with many of these being involved in cognitive control ^50,51^ as well as higher level semantic processing. ^52,53,54,55^ While FLE can derive from different structural causes, a common substrate is focal cortical dysplasia ^56^ (FCD) Type II. Many previous neuroimaging studies in FCD have focused on MRI-based lesion detection ^57^ and modelling, ^56,58^ with only few studies investigating structural and functional alterations beyond the primary substrate. ^56,59^ Similarly, behavioural investigations focused on attention and executive functions, with reduced emphasis and relatively mixed findings on semantic cognition and memory impairment. ^31–33,60^ Interestingly, some studies in FLE patients report post-surgical deficits in free recall ^60,61^ and autobiographical episodic memory, ^33^ while semantic knowledge remains fairly intact. ^33^ Recognition deficits are only observed in a minority of FLE patients, and this group have been shown to often recruit contralateral frontal regions during episodic encoding tasks, suggesting compensatory mechanisms. ^62^ Similarly, semantic fluency impairments appear milder when structured cues are provided, ^63^ suggesting retrieval difficulties rather than a primary degradation of semantic knowledge. ^63^

Individual task-based and rsFC studies in epilepsy, ^30,64–73^ despite showing robust activation or connectivity differences relative to healthy controls, have shown inconsistent relationships with clinical variables. ^30,70,74–78^ Importantly, despite advances in neuroimaging, brain activation and connectivity differences in TLE and FLE have rarely been systematically assessed within a unified framework across cognitive states. A recent task-based fMRI study from our group and collaborators compared TLE and FLE on language and working memory tasks. ^79^ This study revealed a combination of common and syndrome-specific alterations, with diverging patterns of activation and deactivation relative to healthy individuals. Crucially, however, this study did not investigate declarative memory functions more generally, and episodic and semantic memory specifically. Structural and network-level properties, ^56,64,66,80,81^ including microstructural indices and degree centrality, capture brain alterations in TLE and FLE and provide complementary insights into how local and global architecture may support task-evoked activity in these syndromes. Studying TLE related to MTL pathology and FLE related to FCD could therefore provide a window into how temporal and frontal lesions integrate with broader cognitive architectures and impact global brain function in health and disease. For example, the relative preservation of certain memory functions in FLE could inform theories of cognitive resilience and compensation. Conversely, the material-specific memory deficits in TLE ^82,83^ may underscore the key role of MTL structures such as the hippocampus in domain-specific aspects of memory. To examine these network-level effects, AFM offers an opportunity to link intrinsic network disruptions to cognitive impairments without relying on active task performance. By leveraging resting-state data to model memory-related signal flow, AFM enables quantification of cognitive dysfunction and identification of network pathways vulnerable to disruption in temporal and frontal lobe disorders.

This study investigated how intrinsic functional organization contributes to declarative memory network organization and studied system-level alterations in TLE and FLE patients. ^21,23^ All participants underwent episodic and semantic retrieval tasks as well as rs-fMRI in the scanner, from which we computed task activation maps and rsFC estimates. Here, we show: *(i)* that AFM reconstructs memory-related activation patterns in healthy controls but exhibits reduced prediction accuracy in both TLE and FLE; *(ii)* that AFM captures behaviorally and clinically relevant variation not consistently reflected by syndrome-specific dysconnectivity and abnormal activity patterns alone; and *(iii)* that functional and structural features impose distinct relationships with activity flow. These findings provide insight into how focal pathology alters activity flow dynamics and suggest that local lesions and broader network perturbations contribute to memory impairments commonly observed in disorders affecting temporal and frontal lobe systems.

## Results

### AFM predicts state and cohort-specific activations

We estimated participant-specific rsFC and task-evoked activation amplitudes during episodic and semantic retrieval (**Supplementary Figure 1)**. AFM uses rsFC to weight activation from source regions and predict activation in each target region. AFM was first applied to healthy controls (HC_1_;*n*=45; see **Methods**) using their individual rsFC and task activations (**Figure 1a**). Participant-level predicted-to-actual activation overlap was high and significant for both episodic (mean *r*=0.78, *t*=30.13, *P*<0.0001) and semantic tasks (mean *r*=0.78, *t*=30.59, *P*<0.0001). The predicted and actual task activation maps also spatially correlated with the meta-analytic activation maps derived from *Neurosynth*, particularly the semantic task maps (*P_SPIN_*<0.01; **Supplementary Figure 2**). To model the emergence of task dysfunctions in patients and to validate the approach in an independent HC_2_ group, we applied the method using each individual’s rsFC and group-level task activations from HC_1_. As such, AFM was able to reliably simulate activation patterns in an individual based on their rsFC profile and normative task-based activations alone. The predicted activations engaged similar regions as the actual task activations for both episodic and semantic memory in all groups (**Supplementary Figure 3**). At the group level, predicted-to-actual task activation correlations for episodic memory were indeed significant in all groups, (HC_2_/TLE/FLE: mean *r*=0.60/0.44/0.43, *t*=14.54/8.73/6.78, all at *P*<0.000001). Findings were equivalent for semantic memory predictions (HC_2_/TLE/FLE: mean *r*=0.57/0.33/0.33, *t*=12.37/6.64/4.89, *P*<0.000001/*P*<0.000001/*P*=0.001; **Figure 1b-c; Supplementary Table 2**). Prediction accuracy was higher during episodic than semantic retrieval (main effect of *task ME_task_* : *F*(1,92)=9.56, *P*=0.003, η*p*^2^=0.10) and higher in HC_2_ than in both TLE (*P*=0.001) and FLE (*P*=0.005; main effect of Group, *ME_group_*: *F*(2,92)=8.86, *P*<0.001, η*p*^2^=0.16), with no difference between TLE and FLE (*P*=0.99). There was no significant interaction (*F*(2,92)=0.81, *P*=0.45, η*p*^2^=0.02). To determine whether differences in resting-state fMRI data quality influenced AFM prediction accuracy, we conducted the mixed repeated-measures ANOVA including mean framewise displacement (FD) and resting-state temporal signal-to-noise ratio (tSNR) as covariates. The *ME_group_* remained significant after adjustment for both covariates (*F*(2,90)=7.03, *P*=0.001, η*p*^2^=0.14), while neither mean FD (*F*(1,90)=0.04, *P*=0.84) nor resting-state tSNR (*F*(1,90)=0.04, *P*=0.84) was significant. Complete resting-state quality-control analyses are provided in the *Supplementary Materials*.

**Figure 1.**
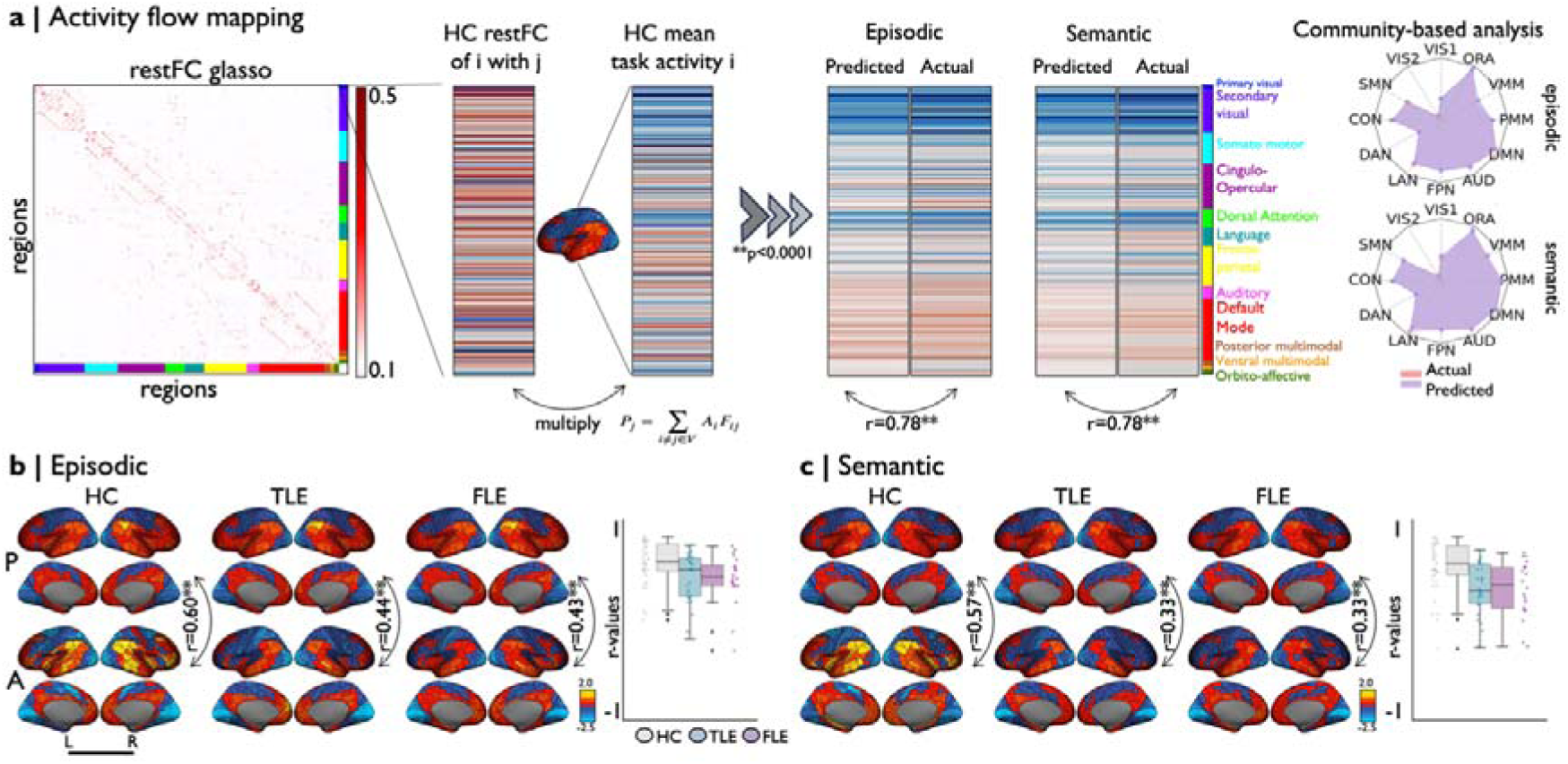
Activity flow mapping (AFM) procedure. **a)** AFM applied to the first group of healthy controls (HC_1_), using their rsFC and task-fMRI activation patterns ^24^, revealed strong correspondence between predicted and actual activations for both tasks. Community-wise analysis revealed task-activation overlap with several large-scale networks^141,142^, such as default mode, posterior and ventral multimodal, orbito-affective, fronto-parietal, and language networks. **b-c)**. Overlap between predicted (P) and actual activations (A), showing significant correspondence across groups, but was reduced in patients. Individual prediction accuracy for HC_2_/TLE/FLE (*n*=42/33/20). Box plots show the median (central line) and interquartile range (box); whiskers extend to the most extreme observations within 1.5 times the interquartile range. Dots represent individual observations. **Abbreviations** | HC=healthy controls, TLE=temporal lobe epilepsy, FLE=frontal lobe epilepsy, VIS1=primary visual, VIS2=secondary visual, SMN=somatomotor, CON=cingulo-opercular, DAN=dorsal attention network, LAN=language, FPN=fronto-parietal network, AUD=auditory, DMN=default mode network, PMM=posterior multimodal, VMM=ventral multimodal, ORA=orbito-affective.

Several analyses verified robustness. First, repeating the analyses in equal-size, age-(*F* (2,57)=0.06, *P*=0.95) and sex-matched (χ*2*(2)=1.67, *P*=0.43) cohorts (*n*=20/group, HC_2_ age=32.50±13.42,F/M=14/6; TLE=32.55±8.89, F/M=12/8; FLE=31.50±10.73, F/M=10/10) yielded comparable results for both episodic (HC_2_/TLE/FLE: mean *r*=0.63/0.46/0.43, *t*=12.02/8.11/6.78, all *P*<0.000001) and semantic tasks (HC_2_/TLE/FLE: mean *r*=0.62/0.29/0.33, *t*=8.14/4.25/4.89, *P*<0.00001/*P*=0.0004/*P*=0.0001). Prediction accuracy remained higher during episodic than semantic retrieval (*ME_task_*: *F*(1,57)=10.31, *P*=0.002, η*p*^2^=0.15) and higher in HC_2_ than in both TLE (*P*=0.005) and FLE (*P*=0.006), with no difference between patient groups (*P*=0.99; *ME_group_ :F*(2,57)=7.04, *P*=0.002, η*p*^2^=0.20; Group × Task interaction: *F*(2,57)=1.43, *P*=0.25, η*p*^2^=0.05). Second, using an independent group activation template from age- and sex-matched participants across all three groups (HC_1_=10, TLE=10, FLE=10) similarly preserved predicted-to-actual correspondence, but remained generally lower in patients (episodic: HC_2_/TLE/FLE mean *r*=0.58/0.48/0.46, *t*=12.45/9.08/6.82, all *P*<0.000001; semantic tasks: (HC_2_/TLE/FLE mean *r*=0.55/0.37/0.36, *t*=10.71/7.38/5.09, *P*<0.000001/*P*<0.000001/*P*=0.00006; **Supplementary Figure 4**). Prediction accuracy again remained higher during episodic than semantic retrieval (*ME_task_*: *F*(1,92)=8.74, *P*=0.004, η*p*^2^=0.09), with an overall group effect (*ME_group_: F*(2,92)=3.65, *P*=0.030, η*p*^2^=0.07). Although HC_2_ showed numerically higher accuracy than both patient groups, *post-hoc* comparisons revealed differences between HC_2_ and TLE (*P*=0.06), HC_2_ and FLE (*P*=0.08) at trend level only. There was no significant Group × Task interaction (*F*(2,92)=0.79, *P*=0.46, η*p*^2^=0.02). Third, replacing the normative HC_1_ template with each cohort’s own average task activation template yielded similar findings (episodic: HC_2_/TLE/FLE: mean *r*=0.61/0.52/0.50, *t*=13.79/9.80/7.57, all at *P*<0.000001; semantic: HC_2_/TLE/FLE: mean *r*=0.59/0.41/0.40, *t*=11.79/9.56/5.75, *P*<0.000001/*P*<0.000001/*P*=0.00002). Prediction accuracy remained higher during episodic than semantic retrieval (*ME_task_*: *F*(1,92)=7.93, *P*=0.006, η*p*^2^=0.08) and differed across groups (*ME_group_ F*(2,92)=4.31, *P*=0.02, η*p*^2^=0.09), with HC_2_ showing higher accuracy than TLE (*P*=0.04), a trend toward higher accuracy than FLE (*P*=0.05) and no differences between TLE and FLE (*P*=0.97; Group × Task interaction (*F*(2,92)=0.76, *P*=0.47, η*p*^2^=0.02). Finally, repeating the analysis across 50 random splits of the 87 HCs (45 participants for the normative template and 42 for validation) showed that the group differences were robust to the choice of healthy-control participants used to derive the normative template. Prediction accuracy remained higher in the held-out HC group than in both TLE and FLE across all 50 splits for both episodic and semantic memory (see **Supplementary Materials** and **Supplementary Figure 5** for details).

The tasks were designed to differentially engage episodic and semantic memory processes, ^29,30^ and despite substantial overlap in their network activations, task-specific patterns were evident, with episodic retrieval preferentially engaging MTL, whereas ATL did not exhibit a comparable episodic enhancement (see **Supplementary Materials**). Across all groups, episodic activations were predicted more accurately than semantic activations, as demonstrated with the main and robustness analyses described above. To assess task specificity, we compared each participant’s predicted activation with group-level actual activations for the same task *vs* the alternate task ^24^ (leave-one-out method; excluding the target participant). The resulting Pearson’s *r* values for each subject were Fisher-*z* transformed and compared using paired-sample *t*-tests. Predicted episodic activations more closely matched actual episodic than semantic activations (HC_2_/TLE/FLE: *t*(41/32/19)=15.87/40.57/14.41, *d*=2.45/7.06/3.22; all *P*<0.000001). In contrast, predicted semantic activations preferentially matched actual semantic activations in HC_2_, but not in both patient groups (HC_2_/TLE/FLE: *t*=21.01/-27.92/-9.53, *d*=3.24/-4.86/-2.13, all *P*<0.000001). When predicting activations in patients, AFM relied solely on individual rsFC and HC_1_ activation vectors. To evaluate cohort specificity, we compared each patient’s predicted activation pattern to their own group’s actual activation ^24^ and to that of the other epilepsy cohort. During the episodic task, predictions were cohort specific in TLE (*t*(32)=2.78, *P*=0.009, *d*=0.48), but not in FLE (*t*(19)=−2.06, *P*=0.05, *d*=-0.46). During the semantic task, TLE predictions more closely resembled the FLE predictions than the TLE activation pattern (*t*(32)=−6.50, *P*<0.000001, *d*=-1.13), whereas FLE predictions preferentially matched the FLE activation pattern (*t*(19)=5.10, *P*=0.00006, *d*=1.14; **Supplementary Table 2**). These results demonstrate that rsFC can transform healthy activation patterns into patient activation profiles while capturing task- and cohort-dependent differences in memory network organization. Finally, local differences indexed by mean absolute errors (MAE) ^84^ between predicted and actual task activity across groups showed higher errors in unimodal regions (episodic/semantic: *t*=-6.77/-6.24, both *P*<0.00001, *d*=0.70/0.64), reflecting localized information flow, compared to transmodal regions involved in more distributed processing ^89^ (**Supplementary Figure 6**).

### Relation to structural and intrinsic functional features

We next examined associations between structural and intrinsic functional brain features and AFM prediction accuracies (**Figure 2, Supplementary Table 2**). Among neocortical structural measures, mean cortical thickness (episodic/semantic: *r*=0.04/0.17, *P_FDR_*=0.74/0.20) and structural connectivity-degree centrality (SC-DC; episodic/semantic: *r*=0.08/0.21, *P_FDR_*=0.43/0.08) were not significantly associated with tasks prediction accuracies, while mean quantitative T1 (qT1) relaxometry, a proxy of cortical myelin and microstructure, ^85,86^ demonstrated positive association with episodic but not semantic prediction accuracy (episodic/semantic: *r*=0.34/0.03, *P_FDR_*=0.002/0.12). Functional features showed consistent negative correlations. Mean functional connectivity-degree centrality (FC-DC), a measure of overall functional integration,^26,87^ was negatively associated with prediction accuracies in both tasks (episodic/semantic: *r*=−0.31/-0.24, *P_FDR_*=0.006/0.02), underscoring the influence of intrinsic network dynamics in task-related processing. Similarly, mean neocortical–hippocampal FC was selectively associated, albeit, at the trend-level, with episodic (*r*=−0.21, *P_FDR_*=0.06), but not semantic (*r*=−0.14, *P_FDR_*=0.19) prediction accuracy across all cohorts. We also examined hippocampal-specific features. Similar to neocortex finding, qT1 was the only structural measure positively associated with episodic but not semantic AFM performance (episodic/semantic: *r*=0.31/0.18, *P_FDR_*=0.006/0.08). In contrast, neither hippocampal thickness (episodic/semantic: *r*=-0.03/0.17, *P_FDR_*=0.78/0.20), nor hippocampal DC (episodic/semantic: *r*=0.10/0.04, *P_FDR_*=0.66/0.73) was related to AFM performance.

**Figure 2.**
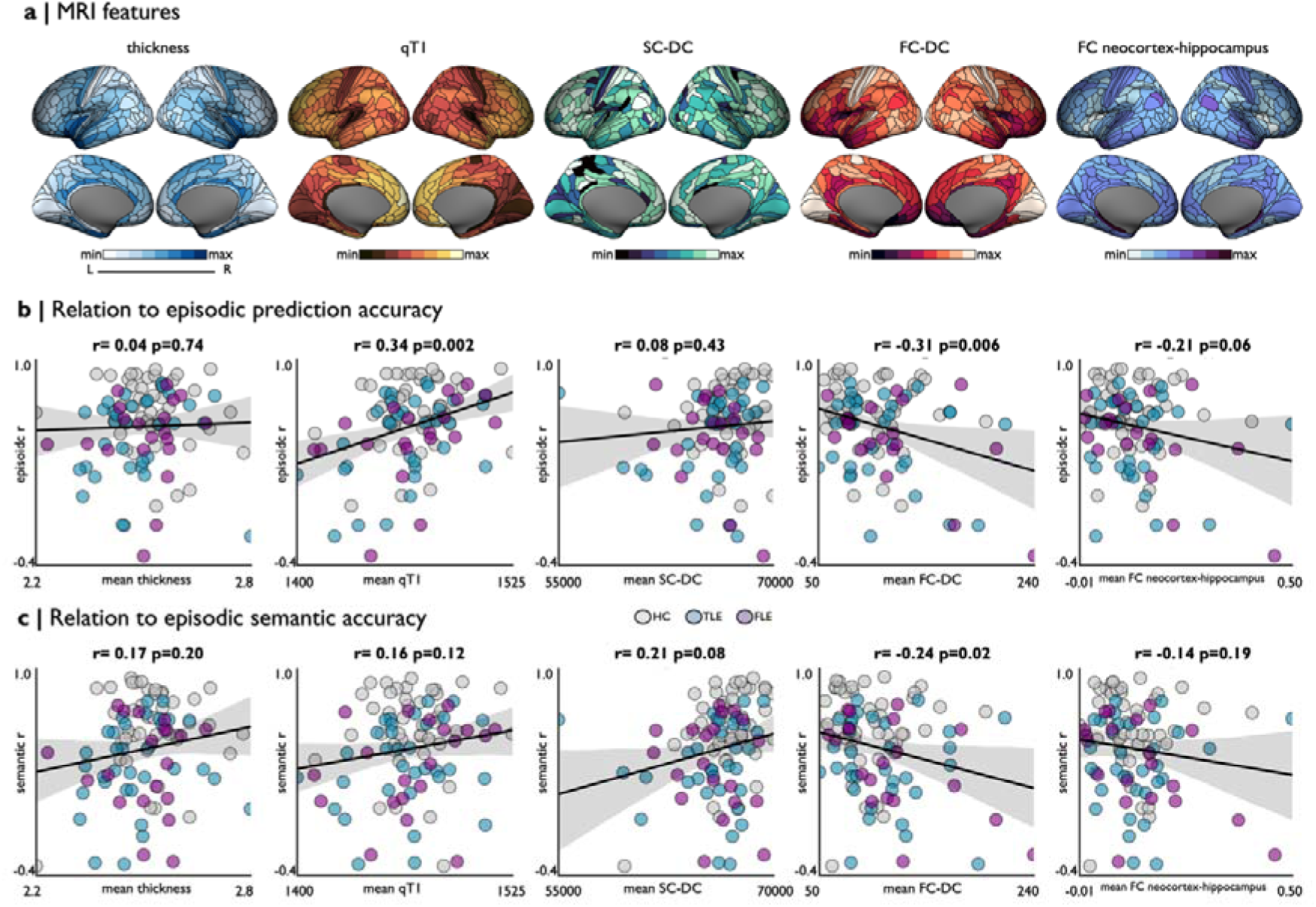
MRI features and activity flow mapping performance. **(a)** Mean structural and functional features from healthy controls. The relationship between each feature, averaged across all nodes, and episodic **(b)** and semantic **(c)** prediction accuracies. Individual data points are colored by group (HC_2_/TLE/FLE; *n*=42/33/20); solid black lines indicate fitted linear regression lines, and shaded regions represent 95% confidence bands. **Abbreviations |** qT1=quantitative T1 image, SC=structural connectivity, DC=degree centrality, FC=functional connectivity, HC=healthy controls (*grey*), TLE=temporal lobe epilepsy (*blue*), FLE=frontal lobe epilepsy (*purple*).

### Behavioural and clinical associations

Finally, we examined behavioural performance (episodic/semantic memory) and EpiTrack scores, a neuropsychological measure of executive functioning commonly used to detect cognitive impairment in epilepsy, ^88^ and its relationship to task-specific prediction accuracy. Behaviourally, episodic memory accuracy differed across groups (*F*(2,92)=15.41, *P*<0.001, η²=0.25). *Post-hoc* Tukey tests showed lower episodic memory accuracy in both TLE and FLE compared with HC_2_ (both *P*<0.001; HC_2_ vs TLE: *d*=1.15; HC_2_ vs FLE: *d*=1.12), with no difference between the patient groups (*P*=0.99, *d*=0.03). Likewise, EpiTrack scores (available for 41 HC_2_, 30 TLE, and 20 FLE participants) differed across groups (*F*(2,88)=12.46, *P*<0.001, η²=0.22), with both patient groups scoring lower than HC_2_ (both *P*<0.001; HC_2_ vs TLE: *d*=1.16; HC_2_ vs FLE: *d*=1.12), but not differing from each other (*P*=0.85, *d*=0.13). In contrast, semantic memory accuracy did not differ across groups (*F*(2,92)=1.07, *P*=0.35, η²=0.02), and *post-hoc* comparisons confirmed no significant differences between any groups (HC_2_ vs TLE: *P*=0.61, *d*=0.24; HC_2_ vs FLE: *P*=0.35, *d*=0.37; TLE vs FLE: *P*=0.85, *d*=0.14).

Descriptive statistics for all behavioural measures and correlations, including FLE-specific sensitivity analyses conducted in light of the smaller sample size, are provided in **Supplementary Table 2 and 3**. Higher AFM prediction accuracies showed a trend-level association with better behavioural performance (episodic/semantic: *r*=0.18/0.21, *P_FDR_*=0.09/0.08). Epitrack scores correlated more significantly with semantic than episodic prediction accuracy (episodic/semantic: *r*=0.08/0.34, *P_FDR_*=0.43/0.002), consistent with semantic retrieval’s reliance on executive control. Disease duration did not differ between patient groups (*t*=-1.97, *P*=0.06). Longer seizure duration negatively correlated to prediction accuracy (episodic/semantic *r*=-0.28/-0.33, *P_FDR_*=0.04/0.03), suggesting disease duration is associated with greater network-level disruptions (**Figure 3).** Because anti-seizure medications (ASMs) may influence functional connectivity and cognitive performance, ^89–91^ we conducted sensitivity analyses adjusting for ASM count and diagnostic group. The overall pattern of associations was broadly consistent with the original analyses, and ASM count was also not significantly associated with AFM prediction accuracy (*see* **Supplementary Materials**).

**Figure 3.**
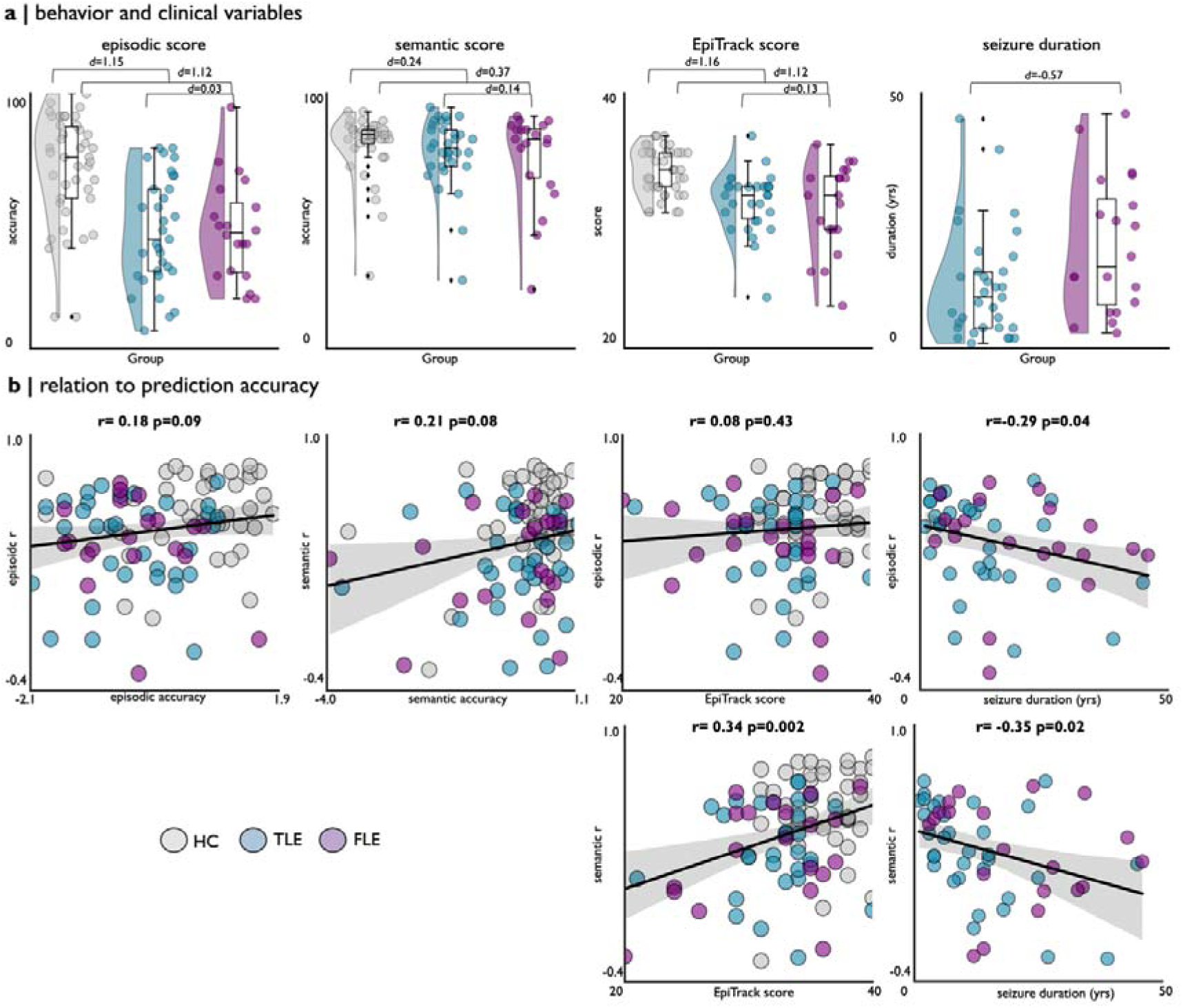
Behavioral and clinical associations. **(a)** In-scanner episodic and semantic scores (%; HC_2_/TLE/FLE (*n*=42/33/20), EpiTrack scores (HC_2_/TLE/FLE (*n*=41/30/20) and seizure duration (TLE/FLE (*n=*33/20). Half-violin plots illustrate the distributions of individual participants, with dots representing individual observations. Box plots show the median (central line) and interquartile range (box); whiskers extend to the most extreme observations within 1.5 times the interquartile range. **(b)** Relation to episodic and semantic prediction accuracies. Individual data points are colored by group; solid black lines indicate fitted linear regression lines, and shaded regions represent 95% confidence bands. **Abbreviations |** HC=healthy controls (grey), TLE=temporal lobe epilepsy (blue), FLE=frontal lobe epilepsy (purple).

Importantly, although clear task- and resting-state abnormalities were present, these conventional measures showed generally more limited associations with clinical variables. An exception was a positive association between semantic task-activation differences and EpiTrack scores in TLE. Task activation abnormalities included reduced activations in higher order networks such as default mode and fronto-parietal networks in both patient groups, alongside marked ipsilateral temporal rsFC disruptions in TLE and ipsilateral frontal/cingulo-opercular rsFC disruptions in FLE. We additionally examined resting-state FC-DC but found no significant parcel-wise group differences or robust associations between mean FC-DC and clinical measures. These findings contrast with the clinical associations observed for AFM prediction accuracy and suggest that integrating intrinsic connectivity with task-activation patterns may capture clinically relevant variation not consistently reflected in conventional measures alone. Complete comparisons are provided in the **Supplementary Materials**.

## Discussion

Understanding how large-scale brain network architecture supports memory and how its disruption leads to cognitive dysfunction is central to network-based models of cognition. Rather than reflecting isolated regional abnormalities, memory deficits in focal epilepsy provide a window into how disruptions of distributed communication pathways alter the emergence of task-evoked cognitive states. Within this framework, AFM provides a computational framework grounded in models of how task-related activity spreads through intrinsic brain networks and how deviations from normative flow constrain memory function. ^74,75^ Unlike many complex models, AFM is grounded in established principles of neural communication that are empirically testable. ^21,23^ In healthy controls, AFM accurately reconstructed episodic and semantic activations, indicating that intrinsic functional architecture scaffolds declarative memory states. Because AFM models activity flow over intrinsic connectivity, prediction accuracy provides an index of disruptions in large-scale activity-flow organization. Consistent with our hypothesis, we found atypical and reduced AFM predictions in both TLE and FLE for both episodic and semantic states, suggesting that intrinsic network disruptions were associated with a reduced ability to generate normative task activations. Both functional and structural MRI features related to variable prediction accuracy, suggesting distinct contributions of dynamic network function *vs* structural integrity to AFM. Importantly, greater mismatch between predicted and actual activations was associated, at the nominal level, with both poorer memory performance and longer disease duration. These associations were generally not detected when task-activation, rsFC alterations and FC-DC are considered separately. Together, these findings indicate that memory impairment in focal epilepsy reflects integrated breakdown of large-scale flow. Even when key memory network nodes may remain structurally intact, atypical network communication may limit task-related activity from being effectively routed through broader declarative memory systems. More broadly, this work advances a network-based perspective on declarative memory by demonstrating how intrinsic brain architecture constrains memory in health and that its disruption gives rise to dysfunction in conditions affecting temporal and frontal lobe systems.

We leveraged AFM to investigate the interplay between intrinsic network alterations, atypical task activations, and cognitive deficits in both TLE and FLE. To our knowledge, this is the first application of AFM to declarative memory, integrating episodic and semantic retrieval within a unified and validated tasks design, ^29,30^ and examining these mechanisms in both healthy and well characterized TLE and FLE cohorts, populations that are difficult to recruit but critical for system-level investigations. This framework allowed us to examine frontal and temporal systems in relation to association *vs* executive control networks. We validated the model in a HC_1_ group, yielding high prediction accuracy across episodic and semantic tasks, consistent with prior work showing that distributed network dynamics shapes task responses. ^19,21,23^ We then extended AFM to TLE and FLE, using each patient’s rsFC and a group-level task activation template from HC_1_ as input. ^24^ Episodic activations were more accurately predicted than semantic activations across groups, a pattern consistent with the more coherent and structurally constrained hippocampal-DMN pathways supporting episodic processing. ^12,13,92^ In contrast, semantic representations are more distributed and heteromodal, ^15,16^ likely making its broader and more diffuse disruptions harder for AFM to predict. Despite significant predicted-to-actual activation overlap, AFM prediction accuracy was consistently lower in patients, suggesting that alterations in intrinsic rsFC may influence the ability of networks to re-instantiate normative task activations. This reduction persisted after robustness tests using mixed-group and cohort-specific activation templates, equalized cohort sizes, and repeated random healthy control splits, indicating that altered rsFC, rather than template source or sample size, contributed to the lower predictions in patients. The FLE cohort encompassed seizure foci and/or lesions across multiple frontal subregions, reflecting the recognized heterogeneity of FLE. Prior work in other conditions has shown that lesions affecting dorsolateral prefrontal cortex might disrupt frontoparietal control processes, whereas lesions involving ventrolateral prefrontal regions may impair semantic control and distributed semantic representation network.^93–95^ Such anatomical variability could therefore contribute to differences in cognitive dysfunction across patients. On the other hand, comparable AFM prediction accuracy in TLE and FLE suggests that the observed network communication deficits were not likely specific to a particular frontal subregion. Importantly, AFM preserved episodic task-specificity, supporting the ability of rsFC to reconstruct task-relevant activation patterns. In contrast, cohort-specificity was task-dependent, indicating that AFM captures both shared and distinct memory aspects of memory network organization, rather than simply reproducing generic task activation patterns. These findings show that models grounded in empirically measured fMRI connectivity can sensitively detect functional alterations across groups.^23^

Distinct structural and functional features contributed to AFM model accuracy. Specifically, higher mean neocortical and hippocampal qT1 were associated with better episodic prediction accuracy, whereas higher FC-DC was associated with lower prediction accuracy in both memory domains. Neocortical-hippocampal FC also showed a negative trend for episodic, but not semantic prediction. Previous studies report that atrophy and microstructural alterations in TLE and FLE co-occur with functional alterations, particularly in mesiotemporal and fronto-limbic regions. ^56,64^ ^3,26,81,85,96–102^ More broadly, anatomical architecture provides the structural scaffold for information flow, while cortical microstructure and intrinsic functional dynamics shape how neural activity propagates through brain networks. ^1,3,103^ In focal epilepsy, pathology can induce structure-function decoupling ^104, 105^ through mechanisms such as diaschisis, ^3,106^ altered excitation-inhibition balance, ^107^ or compensatory reweighting of alternative pathways. ^2^ As a result, regions with structural alterations may preserve or reorganize connectivity, whereas others show disrupted communication despite intact structure. Functional integrity mediates the relationship between structural damage and memory asymmetry in TLE, ^108^ and is often more closely related to cognitive impairment than structural measures alone in TLE and FLE. ^109^ ^110^ These findings suggest that AFM accuracy reflects an interaction between microstructure and functional network dynamics rather than gross morphometric damage or global structural network topology, consistent with the evidence that qT1 detects microstructural abnormalities beyond cortical atrophy and supports memory function. ^85^ ^111^ Within this framework, the negative associations with functional metrics may reflect pathological network reorganization rather than efficient network integration. Although resting-state connectivity typically supports coordinated communication across distributed brain networks, in epilepsy increased FC or greater network centrality may also relate to maladaptive synchronization, aberrant coupling, or compensatory network reorganization.^112^ Finally, the selective association between episodic accuracy and neocortico-hippocampal FC underscores a key role of hippocampal-cortical integration in episodic memory systems. ^13,30,92,113,114^

Beyond group-level network disruptions, AFM prediction accuracy captured inter-individual variation in memory function, indexing how well intrinsic connectivity constrained task-evoked memory states. While associations with memory performance in both domains were attenuated after correction for multiple comparisons, EpiTrack scores were more strongly associated with semantic than episodic prediction accuracy, consistent with semantic retrieval’s reliance on executive and control processes. ^15,53,115^ Longer disease duration was associated to poorer AFM accuracy suggesting that chronic epileptic activity may destabilize network communication. ^116,117^ Crucially, these relationships emerged only and consistently, when rsFC and task activations were modeled jointly. Given that ASMs can alter task-related functional MRI measures, ^89^ ^91^ we also examined associations between the number of ASMs and AFM prediction accuracy, but did not observe robust effects (see **Supplementary Materials** for details). Note, however, heterogeneous medication classes and dosages, as well as sometimes changes in these throughout patient’s journey, precluded systematic assessment of medication-specific effects. Several studies have suggested that conventional brain-behavior correlations may require highly-powered samples to detect robust effects. ^118,119^ On the other hand, studies leveraging well-characterized clinical populations, high-quality phenotyping, and sensitive analytical methods can detect meaningful associations with far smaller samples, ^120–124^ consistent with the robust AFM-behavior relationships observed in this study. The relatively small FLE cohort (*n*=20) limits the precision of subgroup-specific estimates. Although LOO analyses preserved the direction of five of the six FLE-specific correlations, the bootstrap confidence intervals included zero for all six associations, underscoring the uncertainty of these estimates and the need for replication in larger cohorts. In healthy brains, rsFC provides a stable scaffold that channels task-related information flow, functioning as a macroscale analogue of synaptic weights that shape how activity is routed during cognition. ^125,126,127^ In epilepsy however, this mapping becomes unstable as networks can exhibit excessive local synchrony, weakened long-distance coupling, ^125,128^ and state-dependent connectivity shifts, all of which can disrupt coherent information routing and degrade the fidelity of rsFC as a predictor of task activation. Concurrently, atypical or compensatory task recruitment introduces task-specific communication routes that are not evident at rest, ^129,130^ leading to additional mismatches between intrinsic connectivity and task demands. By integrating both resting-state coupling and task-evoked signals, AFM indexes how well intrinsic networks can support the flow of task-relevant activity. This joint influence, rather than rest or task fMRI metrics alone, captures impairments in signal routing and network integration that align with disease progression. The association between AFM accuracy and longer seizure duration underscores alterations in large-scale communication pathways in longer-standing epilepsy, supporting further investigation of whether early seizure control may help preserve cognitive function. ^131^

While our primary aim was to characterize how resting-state and task-evoked disruptions contribute to memory impairments in conditions affecting temporal and frontal systems, we also identified informative differences between episodic and semantic memory that enrich theoretical models of human memory, highlighting epilepsy’s value as a disease model. Behaviorally, we observed episodic memory impairments but no significant impairment in semantic memory performance in TLE, ^28–30,76^ and importantly this pattern extended to FLE, where semantic behaviour also remained intact despite measurable neural disruption. This reflects a partial dissociation between neural and behavioural profiles across memory systems based on the two disease models studied here. In TLE, while both episodic and semantic networks showed reduced task activations, only episodic performance was impaired. This is consistent with accounts ^43,44,132^ postulating that episodic retrieval depends on hippocampal-neocortical integration and is therefore more susceptible to medial temporal disruption. In contrast, semantic memory was not significantly impaired in either patient group despite altered activations, likely echoing its support from a distributed and redundantly organized anterior temporal-heteromodal network, ^15,16^ which enables functional resilience when local regions are affected. In FLE, the absence of significant semantic behavioural impairment despite altered frontal and temporal recruitment suggests compensatory engagement of contralateral control regions, ^63,133^ whereas episodic performance was impaired despite relatively preserved neural activation patterns, likely due to network-level integration failures between the DMN and fronto-parietal systems rather than focal regional dysfunction. ^12,13,71,73,92^ These findings suggest that reduced AFM prediction accuracy may parallel distinct forms of altered network organization across cognitive systems, including adaptive reorganization when behaviour is preserved. Thus, episodic memory emerges as a shared point of vulnerability across syndromes, whereas semantic memory showed relative behavioural resilience even in the presence of neural disruption due to its distributed representational architecture.

Our work underscores the value of generative network neuroscience approaches and the use of human disease models for understanding how intrinsic brain architecture supports and constrains cognition. By integrating rsFC and task-evoked responses, AFM allowed us to move beyond describing regional hypoactivation to identifying how network disruptions plausibly alter the flow of task-relevant signals. Our findings show that memory impairment in focal epilepsy is not solely determined by the location of the core pathology and presumed seizure-generating region, but by the degree to which epilepsy syndromes impact large-scale network communication. More broadly, resting-state-based network modelling offers a principled approach for characterizing how intrinsic brain architecture may constrain cognitive state expression and how disruption of large-scale communication may give rise to memory dysfunction in temporal and frontal lobe systems.

## Materials and Methods

### Participants

Participants were recruited between May 2018 and April 2026 and provided informed consent. We studied 33 TLE patients (left/right=18/15, age=36.64±11.99 years; F/M: 15/18), 20 FLE patients (left/right=10/10, 31.5±10.73 years; F/M: 10/10), and 87 healthy controls (HC; age=32.54±9.08 years, F/M: 40/47). Demographic information and clinical details in patients were collected by interviewing them and their family members. All patients were carefully selected by experts (TA, NB, AB) based on evaluation of the medical history, neurological examination, seizure semiology, video-EEG, and clinical neuroimaging. Clinical information included: *(i)* mean age at seizure onset (TLE=23.58±13.92 years, range=0.5-60 years; FLE=11.73±6.16, range=0.8-22 years); *(ii)* mean epilepsy duration (TLE=12.64±10.60 years, range=1-45 years; FLE=19.35±13.23, range=3-46 years); *(iii)* history of childhood febrile convulsion (TLE=6, FLE=2); *(iv)* family history of epilepsy (TLE=5, FLE=6); *(v)* anti-seizure medication (ASMs; range for all patients=0-5) with differing dosage. Based on quantitative hippocampal MRI volumetry, ^134^ 18/33 (55%) TLE and 4/20 FLE patients (20%) showed marked hippocampal atrophy ipsilateral to the focus (*i.e.,* absolute ipsilateral volume or ipsilateral–contralateral asymmetry *z*-score >1.5 of the corresponding mean of healthy controls). See **Supplementary Table 1** for more detailed clinical characteristics. All MRI and behavioral data used in the analyses were acquired during the same study session and before resective surgery and/or thermocoagulation. At the time of the study, 11 TLE patients had undergone resective surgery, and one underwent thermocoagulation. Seizure outcome was assessed using Engel’s ^135^ modified classification, with an average follow-up of 27±22.4 months. Ten TLE patients achieved complete seizure freedom (Engel-I) post-surgery, while one had recurring seizures. Histopathological analysis of ten specimens showed mesiotemporal/hippocampal sclerosis in five patients, mesiotemporal ganglioglioma in one, cavernoma in one, gliosis in three, and mild dysplasia in the mesiotemporal structures in one. Four FLE patients underwent resective surgery (2/4 preceded by thermocoagulation), and two had thermocoagulation. Seizure outcomes, with an average follow-up of 27±20.7 months, showed that three FLE patients (75%) achieved complete seizure freedom (Engel-I), while one had recurring seizures. Histopathological analysis of four specimens confirmed FCD type II. Healthy controls were aged 18-65 years, with no history of neurological or psychiatric disorders, substance abuse, brain injury or surgery. There were no significant age differences across HC, TLE and FLE groups (one-way analysis of variance ANOVA: *F*(2,137)=2.38, *P*=0.10. There were also no significant differences in sex distribution across groups (χ*2*(2)=0.12, *P*=0.94). The study was approved by the Montreal Neurological Institute and Hospital Research Ethics Board (MP-37-2018-3469/ MEO-37-2023-4738) and conducted per the Declaration of Helsinki.

### MRI acquisition

MRI data were acquired at the McConnell Brain Imaging Centre of the Montreal Neurological Institute on a 3T Siemens Magnetom Prisma-Fit with a 64-channel phased array head coil. Controls and patients completed the same research protocol which included: *(i)* T_1_-weighted MRI with a 3D-magnetization-prepared rapid gradient-echo sequence (MPRAGE; 0.8 mm isovoxels, matrix=320×320, 224 sagittal slices, repetition time (TR)=2300 ms, echo time (TE)=3.14 ms, inversion time (TI)=900 ms, flip angle=9°, iPAT=2, partial Fourier=6/8, field of view (FOV)=256mm×256 mm); *(ii)* T_1_ relaxometry using a 3D-MP2RAGE sequence (0.8 mm isovoxels, 240 sagittal slices, TR=5000 ms, TE=2.9 ms, TI_1_=940 ms, TI_2_=2830 ms, flip angle_1_=4°, flip angle_2_=5°, FOV=256mm×256 mm); *(iii*) diffusion-weighted imaging (DWI) using a two-dimensional spin-echo echo-planar imaging sequence with multiband acceleration (1.6 mm isovoxels, three diffusion-weighting shells with b values of 300, 700, and 2000 s/mm^2^ acquired with 10, 40 and 90 diffusion directions, respectively, TR=3500 ms, TE=64.40 ms, flip angle=90°, refocusing flip angle=180°, FOV=224mm×224 mm, slice thickness=1.6 mm, multiband factor=3, echo spacing=0.76 ms); *(iv)* resting-state fMRI and; *(v)* episodic and semantic task-fMRI with a 2D/blood oxygenation level-dependent echo-planar imaging sequence (3.0 mm isovoxels, matrix=80×80, 48 slices oriented to AC-PC-30°, TR=600 ms, TE=30 ms, flip angle=50°, FOV=240mm×240 mm, slice thickness=3 mm, multi-band factor=6, echo spacing=0.54 ms). The resting-state fMRI session lasted ∼6 min, during which participants fixated on a central grey cross and allowed their minds to wander. The event-related episodic and semantic retrieval fMRI tasks also lasted ∼6 min. Each task consisted of 56 trials, matched for low-level structure by using symbolic stimuli and a three-alternative forced-choice design. In the episodic task, participants were asked to view and memorize paired images of objects, and a retrieval phase was administered 10 mins later. In the latter phase, for each trial, participants were shown a cue object at the top of the screen and three choices at the bottom and were asked to identify the object paired with the cue during encoding. Data analysis focused only on the retrieval phase. The semantic task consisted of a retrieval phase, where participants saw a cue object and three choices, and were asked to identify the object most conceptually related to the prime (**Supplementary Figure 1a, b**). Task details are reported elsewhere. ^29,30^

### MRI processing

MRI data were preprocessed with *micapipe* v.0.2.3 (https://github.com/MICA-MNI/micapipe), an open-access multimodal preprocessing and data co-registration software. Processing steps are detailed elsewhere. ^136^ Briefly, T1-weighted images were de-obliqued and reoriented to ensure consistent orientation across all datasets. Subsequently, linear co-registration, intensity non-uniformity correction, and skull stripping were applied. DWI data underwent denoising, motion and eddy current correction, and bias field non-uniformity correction. Resting-state and task-fMRI data underwent preprocessing using *FSL* ^137^ v.6.0.2, *ANTs* ^138^ v.2.3.4, and *AFNI* ^139^ v.23.1.09. This process included removal of the initial five TRs, reorientation of the images, motion and distortion correction using AP-PA blip field maps, high-pass filtering (>0.01 Hz), *MELODIC* decomposition, non-linear co-registration to the T1-weighted scans, nuisance signal removal via ICA-based X-noiseifier, ^140^ registration to native cortical surfaces, surface-based registration to the Conte69 template with 32k surface points per hemisphere, statistical regression of motion parameters, and resampling to the Human Connectome Project multimodal parcellation atlas ^141^ ^142^ (360 nodes, Glasser parcellation).

### Resting-state functional connectivity estimation

For rsFC estimation, we followed the AFM framework, which commonly employs regularized graphical lasso ^143^ that yields more robust and interpretable estimates of direct functional relationships. Specifically, we computed the rsFC using the *GGLasso* Python package ^144^ and resulted in a 360×360 connectome. This method applies partial correlation with an L1 penalty to control model complexity. ^143,145,146^ The L1 penalty is added during the computation of the precision matrix, which is then transformed into the partial correlation matrix. The penalty term, (**λ_1_** Σ*_j≠k_* |P*_jk_* |), is proportional to the sum of the absolute values of the off-diagonal elements in the estimated precision matrix (P), with the regularization strength controlled by the hyperparameter 1. This L1 regularization encourages sparsity by driving less informative coefficients to zero, effectively improving the reliability of rsFC estimates. Following the default ActFlow toolbox v.0.3.2 implementation (https://github.com/ColeLab/ActflowToolbox), the regularization parameter (**λ_1_**) was selected independently and automatically for each participant using 10-fold cross-validation. Candidate λ values comprised a logarithmically spaced sequence ranging from 0.316 to 0.001 (10 · to 10 ³, in 0.1 log increments), with the optimal value chosen by minimizing the cross-validated negative log-likelihood. ^143^ We used this default, recommended Actflow procedure, ^143^ which provides a data-driven estimate of the regularization strength for each participant rather than relying on an arbitrary fixed parameter.

### Task-fMRI activation estimation

Participant- and task-fMRI specific activation amplitudes were estimated using a GLM convolved with the SPM canonical hemodynamic response function (including derivatives and dispersion) in *nilearn* v.0.11.2. All retrieval trials, regardless of accuracy, were included and modelled as events. Region-specific contrast estimates during the retrieval phase were computed with six motion parameters as confound regressors. The resulting β-amplitude estimates (360 regions × 2 tasks × 140 participants) were used in all analyses involving fMRI task activations.

### Activity flow mapping

The AFM technique was applied to quantify the relationship between rsFC and task-activation patterns, following established protocols and prior recommendations. ^21,23^ Healthy controls were randomly partitioned into a training cohort (HC_1_, *n*=45, age=32.44±6.75 years, F/M: 23/22) and an independent validation cohort (HC_2_, *n*=42, age=32.64±11.13 years, F/M: 17/25 while ensuring that both subgroups were comparable in age and sex and remained demographically matched to the patient cohorts. We first applied the technique to HC_1_ using their own rsFC and task-activation amplitudes. ^21,24^ The AFM technique predicts the activation of a target brain region *P_j_* by combining the task-related activation amplitudes *A_i_* from source regions *i* and their rsFC *F_ij_* with the target region. Specifically, the predicted activation *P_j_* is calculated as the weighted sum of the activation values *A_i_* from all regions *i* except the target region *j*, with the weights being the rsFC *F_ij_*between source and target regions. Mathematically, this is expressed as:

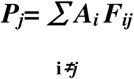

AFM predictions were benchmarked as the overlap between predicted and actual activations across all ROIs. ^21,24^ Specifically, participant-level significance was determined using a random effects model where the overlap correlation (*r* value) for each participant was Fisher-*z* transformed and tested against zero using a one-sample *t*-test. We then repeated this analysis in a second subgroup of healthy controls (HC_2_, *n*=42), and in the TLE and FLE. Notably, models were only trained using data from HC_1_ when predicting task activity in HC_2_, TLE, and FLE. In this modified AFM model, the task activation was sourced from HC_1_ (*i.e.,* group-averaged β-weights), while the rsFC were derived from each HC_2,_ TLE and FLE participant. This adaptation, also used elsewhere, ^24^ allows the procedure to be applied even when memory task-fMRI is unavailable. While our analyses primarily focused on global similarity, mean absolute errors ^84^ (MAE) between predicted and actual task-evoked activity for each memory state and all participants were also computed to show local patterns of errors, where lower MAE indicated better prediction of the target region’s activity.

### MRI features and task-activity flow

Several functional and structural MRI-derived features were extracted to provide complementary information on intrinsic brain organization and assess how these properties reflected AFM predictions. As in previous work from our group ^66,96^ and others, ^147^ neocortical thickness was measured as the Euclidean distance between corresponding pial and white matter vertices, and we used mid-thickness surfaces to sample co-registered quantitative T1 relaxometry data as an index of cortical microstructure. ^80,148,149^ Neocortical degree centrality from rsFC (FC-DC) and structural connectome (SC-DC) was calculated by summing all weighted cortico-cortical connections for each brain region. To ensure spatial alignment, all neocortical features were registered to the Conte69 surface template and resampled using Glasser parcellation. Hippocampal thickness, qT1 values and DC were also computed from hippocampal surface meshes extracted using *HippUnfold* ^134^ v.1.0.3 (https://hippunfold.readthedocs.io/). However, hippocampal SC was excluded because the complex laminar architecture and rapidly changing fiber orientations of the hippocampus limit reliable reconstruction with current *in vivo* diffusion MRI tractography. ^150,151^ Finally, FC between bilateral neocortex and hippocampus was assessed via correlations of timeseries from all neocortical and hippocampal regions and averaged across the hippocampal regions.

### Neurosynth maps

We examined the spatial alignment between actual task activation maps, predicted task activation maps, and task-fMRI patterns from *Neurosynth* ^152^ (https://neurosynth.org/). The latter is a platform for large-scale *ad hoc* meta-analysis of task-based fMRI data. From *Neurosynth*, we retrieved task activation maps for the terms “episodic memory” (332 studies) and “semantic memory” (1031 studies). Maps were registered to the Conte69 surface template and resampled using Glasser parcellation. ^141^ ^142^ Given the larger number of studies contributing to semantic map, differences in statistical power should be considered when interpreting spatial alignment.

### Statistical analysis

To compare AFM prediction accuracy across groups and tasks, we performed a mixed repeated-measures ANOVA with *group* as the between-subject factor and *task* as the within-subject factor. We also evaluated the relationship of AFM’s prediction accuracy and mean functional and structural features, seizure duration, task performance and EpiTrack scores, ^88^ a clinical index of executive functions and attention deficits in epilepsy, assessing processing speed, cognitive flexibility, response inhibition, working memory, planning and verbal fluency.^88^ For the correlation analyses, false discovery rate (FDR) correction was applied separately for each MRI, behavioral, and clinical measures across the episodic and semantic prediction tasks (*P_FDR_*<0.05). Additional exploratory analyses examining hippocampal volume, ASMs and AFM prediction accuracy were also conducted. Effect sizes are reported as partial eta-squared (_η_*p*^2^) for ANOVA models, Pearson’s *r* for correlation analyses, and *Cohen’s dz* for paired-sample *t*-tests. We then conducted a secondary analyses to compare actual task activation between patients and controls using surface-based linear models implemented in *Brainstat* ^153^ v.0.4.2 (https://brainstat.readthedocs.io), controlling for age and sex. Region-wise effects were corrected for multiple comparison using FDR correction (*P_FDR_*<0.05). Group differences in rsFC (360×360) was also assessed and corrected for multiple comparisons at the family-wise error level (*P_FWE_*<0.05), given the large number of correlated edges. Prior to the surface-based group comparisons of task activation and rsFC only, patients’ data were left-right flipped such that the seizure-generating hemisphere was represented on the left, and the contralateral hemisphere was on the right. ^64,148,154,155^ This procedure was performed solely to align the epileptogenic hemisphere across patients for group-level spatial analyses. All AFM prediction accuracy estimates were performed on the original unflipped data.

## Data Availability

Our de-identified healthy control cohort comprises a subset of participants from the MICA-MICs ^156^ ^157^ dataset, which is openly available through Open Science Framework (OSF, https://osf.io/j532r) and on the Canadian Open Neuroscience Platform data portal (https://portal.conp.ca/dataset?id=projects/mica-mics). The raw patient imaging data are not publicly available, and there are currently no plans to release them. However, the derivative preprocessed fMRI data, AFM prediction outputs and Source Data file ^158^ are available on OSF (https://osf.io/wgzk2/).

## Code Availability

Our AFM implementation used in this study was adapted from the AFM toolbox v.0.3.2 (https://github.com/ColeLab/ActflowToolbox) and the AgingActflow_release repository (https://github.com/ColeLab/AgingActflow_release). ^24^ Custom scripts developed for this study are also available in OSF (https://osf.io/wgzk2/). ^158^

## Acknowledgements

We would like to thank all participants who generously contributed to this study and acknowledge Ilana Ruth Leppert, Ronald Lopez, David Costa, Soheil Quchani, and Michael Ferreira for assistance with data collection. We dedicate this work to the late Jonathan Smallwood for his valuable contributions to this research and to the first author’s PhD journey.

## Author contributions

Conception, design, data processing, analysis and manuscript preparation: DGC, BCB. Conception: DGC, BCB, EJ, LC, SA, AJB, NS. Data analysis: DGC, JD, KX; Participant recruitment: TA, RP; Patient selection: TA, AB, NB; Data acquisition: AN, ES, JR; Data processing: DGC, RRC, ES; All authors provided feedback and approved the final manuscript.

