## Supplementary Materials for "Memory network activity flow failures in temporal and frontal lobe epilepsy"

**RUNNING TITLE:**

Disrupted memory activity flow in focal epilepsies

**Task Validation**

Although behavioral results suggested a distinction between episodic and semantic memory, it remained unclear whether these differences were accompanied by distinguishable neural activation patterns. We therefore conducted a neural-level manipulation check in the healthy control group (*n*=87) to examine whether the two retrieval conditions elicited distinguishable activation patterns in canonical memory-related regions. Regions-of-interest (ROI) were defined a-priori using the Glasser 360-area parcellation. [1,2] The bilateral mesiotemporal lobe (MTL) ROI comprised hippocampus, presubiculum, entorhinal cortex, perirhinal cortex, parahippocampal Areas 1–3 (PHA1-3), while the bilateral anterior temporal lobe (ATL) ROI comprised area TE1/TE2 anterior, TG dorsal and TG ventral. These regions were selected based on established evidence implicating the MTL in episodic memory and the ATL in semantic/conceptual processing. [3-8] At the whole-brain level, a second-level GLM identified greater activation during episodic than semantic retrieval ($P_{FDR}$<0.05) in medial temporal regions (left hippocampus, bilateral PHA3, right PHA1 and PHA2), medial frontal/cingulate regions (right dorsal and ventral area 24d), and posterior association regions (left VMV2 and right POS1). Greater activation during semantic than episodic retrieval ($P_{FDR}$<0.05) was observed primarily in prefrontal regions (right s6-8, 8BL, 8BM, 9p, p47r, and 8Av) and parietal association cortex (right PFm). We then tested the predicted MTL–ATL dissociation. Episodic retrieval produced greater activation than semantic retrieval in the MTL (mean episodic=0.20, mean semantic=−0.25; Δ= 0.45; $t(86)$=3.71, $P_{FDR}$<0.001, $dz$=0.40, 95% *CI* [0.21, 0.69]), whereas the ATL contrast was not significant, despite nominally higher activations during semantic retrieval (mean episodic=0.13, mean semantic=0.21; Δ=−0.071; $t(86)$=−0.60, $P_{FDR}$=0.55, $dz$=−0.06, 95% *CI* [−0.30, 0.16]). These findings were also supported by a significant Region × Condition interaction (Δ=0.52; $t(86)$=5.14, $P_{FDR}$< 0.001, $dz$=0.55, 95% *CI* [0.32, 0.72]), indicating that the episodic–semantic difference was significantly stronger in the MTL than ATL. Thus, the two conditions showed a statistically distinguishable neural profile, with episodic retrieval preferentially engaging the MTL, and subtle, nonsignificant findings in the ATL that nominally pointed towards a greater implication in semantic retrieval. These findings are overall consistent with a relative specialization within an overlapping memory architecture.[9-12]

**Resting-state dysconnectivity and task-evoked dysfunction**

Throughout this study, the terms task dysfunction and dysconnectivity refer to statistically significant group differences in task-evoked activation and resting-state functional connectivity, respectively, relative to healthy controls. In our HCs, episodic and semantic tasks engaged widespread medio-lateral temporal and posterior-parietal regions, extending to larger fronto-parietal (FPN) and default mode networks (DMN). Similar but less extensive activation patterns were observed in TLE and FLE patients, as confirmed by community-wise analysis across both memory states (**Supplementary Figure 1**). Compared to $HC_2$, TLE patients showed reduced episodic task activations, particularly in bilateral temporal, cingulo-opercular, latero-frontal, and visual areas ($P_{FDR}$<0.05, $d$=0.74). On the other hand, FLE patients exhibited less pronounced episodic reductions, mainly in fronto-parietal regions ($P_{FDR}$<0.05, $d$=1.04). For semantic memory, TLE patients showed decreased activation in medio-temporal, bilateral latero-frontal, cingulo-opercular, and visual areas ($P_{FDR}$<0.05, $d$=0.78), while FLE patients demonstrated similar reductions in bilateral temporal and medio-lateral frontal regions ($P_{FDR}$<0.05, d=0.86), relative to $HC_2$. At rest, widespread rsFC disruptions were observed in both groups compared to controls. In TLE, dysconnectivity was most pronounced in bilateral DMN, visual areas and, particularly in the ipsilateral temporal region. FLE patients exhibited comparable bilateral DMN-related dysconnectivity and strong disruptions in ipsilateral fronto-parietal and cingulo-opercular regions (**Supplementary Figure 1** and **Table 4**). These patterns align with known syndrome-specific vulnerabilities. In TLE, episodic and semantic tasks functional alterations overlapped with marked rsFC disruptions, [13-15] reflecting decreased connectivity around the core pathological substrate [16-19] and increased network segregation, [17] leading to a less integrated brain organization. [20,21] These disturbances likely undermines memory processes and have been associated with epileptic activity [22,23] and disease duration . [24,25] In FLE, relatively preserved episodic activation patterns likely reflect the hippocampus-dependent nature of episodic memory, typically spared in FLE [3,4,26] (*i.e.,* only 4/17 FLE patients had hippocampal atrophy). In contrast, semantic activations showed broader reductions [6,27,28] and parallel rsFC disruptions consistent with prior studies, [29-31,32] indicating reduced integration between subnetworks and increased interhemispheric short-range coupling. [29,32,33]

**Comparison of AFM with conventional task-activation and resting-state measures**
To compare the clinical associations of AFM with those of conventional approaches, we examined associations between clinical measures and task-activation differences, resting-state functional connectivity (rsFC) abnormalities, and functional connectivity degree centrality (FC-DC). Episodic task activation in regions showing significant differences between patients and controls was not associated with in-scanner episodic (TLE/FLE: $r$=0.09/0.12, $P_{FDR}$=0.47/0.36), EpiTrack (TLE/FLE: $r$=0.18/0.04, $P_{FDR}$=0.14/0.74) or disease duration (TLE and FLE: $r$=-0.13, $P_{FDR}$=0.37). Similarly, semantic task activation was not correlated with in-scanner semantic (TLE/FLE: $r$=-0.06/-0.01, $P_{FDR}$=0.62/0.92), although it was positively associated with EpiTrack scores in TLE only (TLE/FLE: $r$=0.29/0.17, $P_{FDR}$=0.02/0.17). No association with disease duration was observed (TLE and FLE: $r$=0.06 $P_{FDR}$=0.67). Likewise, rsFC dysfunctions were unrelated to episodic (TLE/FLE: $r$=0.15/0.25, $P_{FDR}$=0.20/0.10) and semantic memory performance (TLE/FLE: $r$=-0.14/0.22, $P_{FDR}$=0.23/0.16), EpiTrack (TLE/FLE: $r$=0.03/0.12, $P_{FDR}$=0.78/0.36) or disease duration (TLE and FLE: $r$=-0.13, $P_{FDR}$=0.37). Although no brain regions showed significant differences resting state degree centrality (FC-DC), mean FC-DC was not associated with episodic (TLE/FLE: $r$=0.03/0.01, $P_{FDR}$=0.76/0.92), semantic performance (TLE/FLE: $r$=0.08/0.14, $P_{FDR}$=0.49/0.29), EpiTrack (TLE/FLE: $r$=-0.10/-0.13, $P_{FDR}$=0.41/0.30) or disease duration (TLE and FLE: $r$=-0.01, $P_{FDR}$=0.94). The effect sizes and clinical associations for AFM and conventional measures are summarized in **Supplementary Table 5**. In contrast, AFM prediction accuracy was associated with executive function and disease duration, while associations with memory performance were evident at the nominal level, highlighting its sensitivity to clinically relevant variation. Thus, AFM provides a complementary mechanistic index of altered activity-flow organization, capturing clinically relevant information beyond what can be inferred from rest- or task-based differences in isolation.

**Sensitivity Analyses**
**AFM Sensitivity to healthy control sampling.** To assess robustness to the $HC_1/HC_2$ split, we repeated the AFM analysis across 50 random splits of the 87 healthy controls (HCs), using random 45 participants to derive normative task-activation template and the remaining 42 for validation. The same HC45-derived template was used to predict task activations in the TLE and FLE groups. Prediction accuracy was consistent across splits in episodic ($HC_2$/TLE/FLE: mean $r$=0.48/0.41/0.40, $SD$=0.03/0.008/0.006; ranges=0.41–0.53/0.39–0.43/0.39–0.42) and semantic memory ($HC_2$/TLE/FLE: mean $r$=0.48/0.31/0.30, $SD$=0.03/0.006/0.005; ranges=0.42–0.56/0.30–0.33/0.30–0.31). The direction of the group effect was stable, such that HC prediction accuracy was higher than TLE and FLE in all 50 splits for both tasks (**Supplementary Figure 5**). Across splits, the mean HC–TLE difference was ~0.06±0.03 for episodic and ~0.16±0.03 for semantic memory, while the corresponding HC–FLE differences were ~0.07±0.03 and ~0.17±0.03, respectively. These results indicate that the lower AFM

prediction accuracy observed in the epilepsy groups was not dependent on the particular $HC_1/HC_2$ split used to derive the normative activation template.

**Head motion and tSNR**. To assess whether differences in resting-state fMRI quality could account for reduced AFM prediction accuracy in epilepsy, we quantified whole-brain temporal signal-to-noise ratio (tSNR) and mean framewise displacement (FD) for all participants. Mean FD was 0.11±0.04/0.15±0.06 /0.15±0.12 mm in $HC_2$,TLE and FLE, respectively. An overall effect of group was observed for mean FD ($F(2,92)=3.60$, $P=0.03$, $\eta^2=0.07$), although *post hoc* Tukey comparisons did not identify significant pairwise group differences after correction for multiple comparisons (all $P>0.05$). Mean whole brain tSNR was 58.65±7.44 in $HC_2$, 54.34±7.77 in TLE, and 50.91±7.72 in FLE. A group difference was observed for tSNR ($F(2,92)=7.61$, $P=0.0009$, $\eta^2=0.142$). Tukey's *post hoc* tests showed that $HC_2$ had higher tSNR than both TLE ($P=0.04$) and FLE ($P=0.0009$), whereas tSNR did not differ between TLE and FLE ($P=0.26$).To determine whether these differences influenced AFM prediction accuracy, we compared the prediction accuracies across all groups using mixed repeated-measures ANOVA including both mean FD and resting-state tSNR as subject-level covariates. The main effect of Group remained significant after adjustment ($F$ (2,90)=7.03, $P=0.001$, $\eta p^2=0.14$), whereas neither mean FD ($F(1,90)=0.04$, $P=0.84$) nor resting-state tSNR ($F(1,90)=0.04$, $P=0.843$) was significant. These findings indicate that the observed group differences in AFM prediction accuracy are unlikely to be explained by differences in head motion or resting-state signal quality in the present sample.

**Seizure laterality.** To assess whether seizure laterality influenced AFM prediction accuracy, we compared Fisher $z$-transformed subject-level prediction accuracies between left- and right-lateralized patients within the TLE and FLE cohorts using independent-samples $t$-tests. No marked laterality effects were observed in either cohort for episodic (TLE/FLE: $t(31/18)=0.43/1.99$, $P_{FDR}=0.70/0.12$, $d=0.15/0.89$) or semantic AFM prediction accuracy (TLE/FLE: $t(31/18)=0.29/0.95$, $P_{FDR}=0.87/0.35$, $d=0.10/0.43$).

**Exploratory Analyses**

We performed exploratory analyses stratifying patients according to the presence (n=18) or absence (n=15) of hippocampal atrophy (HA, i.e., absolute ipsilateral volume or ipsilateral–contralateral asymmetry z-score >1.5 of the corresponding mean of healthy controls).). Associations between hippocampal volume and AFM prediction accuracy showed qualitatively similar patterns across both subgroups, and none of the correlations survived correction for multiple comparisons (**Supplementary Table 2**). Leave-one-out (LOO) analyses yielded similar correlation estimates, suggesting that the observed associations were not driven by individual participants. Nevertheless, these subgroup analyses should be interpreted cautiously given the modest sample sizes. To assess the sensitivity of FLE-specific clinical and behavioral correlations to individual observations, we performed LOO analyses and bootstrap resampling (10,000 iterations). The direction of five of the six correlations was preserved across LOO iterations, whereas the association between episodic AFM accuracy and episodic memory scores varied from negative to near zero. All six bootstrap 95% confidence intervals included zero, indicating substantial uncertainty in the FLE-specific correlation estimates. These findings should thus be interpreted cautiously given the relatively small FLE sample size ($n=20$; **Supplementary Table 3**). Given the potential effects of anti-seizure medications (ASMs) on brain connectivity and cognition, [34-36] we conducted sensitivity analyses to assess whether the associations between AFM prediction accuracy and behavioral or clinical measures were robust to adjustment for diagnostic group and ASM count. The overall direction of associations was generally consistent with the original analyses (see **Results;** episodic memory

(*B*=0.005, *P*=0.85), semantic memory (*B*=0.05, *P*=0.08), EpiTrack–episodic accuracy (*B*=−0.004, *P*=0.589), EpiTrack–semantic accuracy (*B*=0.01, *P*=0.06), seizure duration–episodic accuracy (*B*=−0.006, *P*=0.05), and seizure duration–semantic accuracy (*B*=−0.008, *P*=0.01). ASM count was also not significantly associated with AFM prediction accuracy for either episodic or semantic predictions in the present cohort (**Supplementary Table 2**). We next examined whether the observed brain-behavior associations were independent of diagnostic group by performing partial correlations controlling for group ($HC_2$/TLE/FLE). Given the modest patient subgroup sizes, partial correlations were used in place of formal within-group analyses. The overall pattern was preserved but attenuated after adjustment, with positive associations remaining for AFM prediction accuracy and semantic memory (partial *r*=0.17, *P*=0.09) and EpiTrack performance (partial *r*=0.19, *P*=0.08).

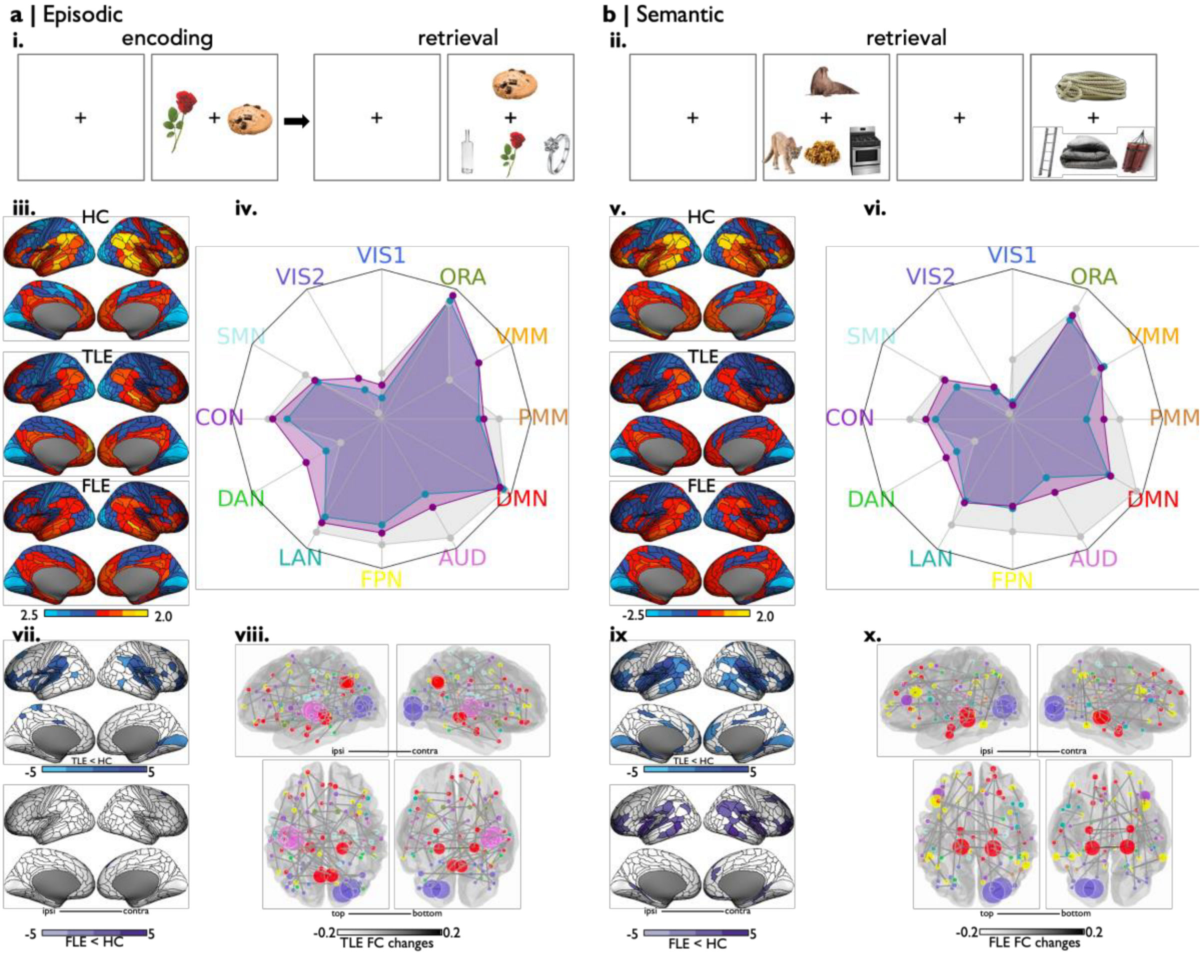


**Supplementary Figure 1. Experimental design, task activation and resting state patterns. (a, b)** Episodic ***(i)*** and semantic ***(ii)*** experimental design. ***(iii,v)*** task activations in healthy, TLE and FLE patients. ***(iv,vi)*** Community-wise analysis showing task-activation maps overlapping with higher-order networks[1,2] (grey=HC, blue=TLE, purple=FLE). ***(vii,ix)*** Regions showing significant task-activation differences between healthy controls and TLE (blue) and FLE (purple). Resting-state functional dysconnectivity in TLE ***(viii)*** and FLE ***(x).*** Node-colours are according to Glasser network assignments. Node size equals to total edge strengths. **Abbreviations** | HC=healthy controls, TLE= temporal lobe epilepsy, FLE=frontal lobe epilepsy, FC=functional connectivity, VIS1=primary visual ,VIS2=secondary visual, SMN=somatomotor, CON=cingulo-opercular, DAN=dorsal attention network, LAN=language, FPN=fronto-parietal network, AUD=auditory, DMN=default mode network, PMM=posterior multimodal, VMM=ventral multimodal, ORA=orbito-affective.

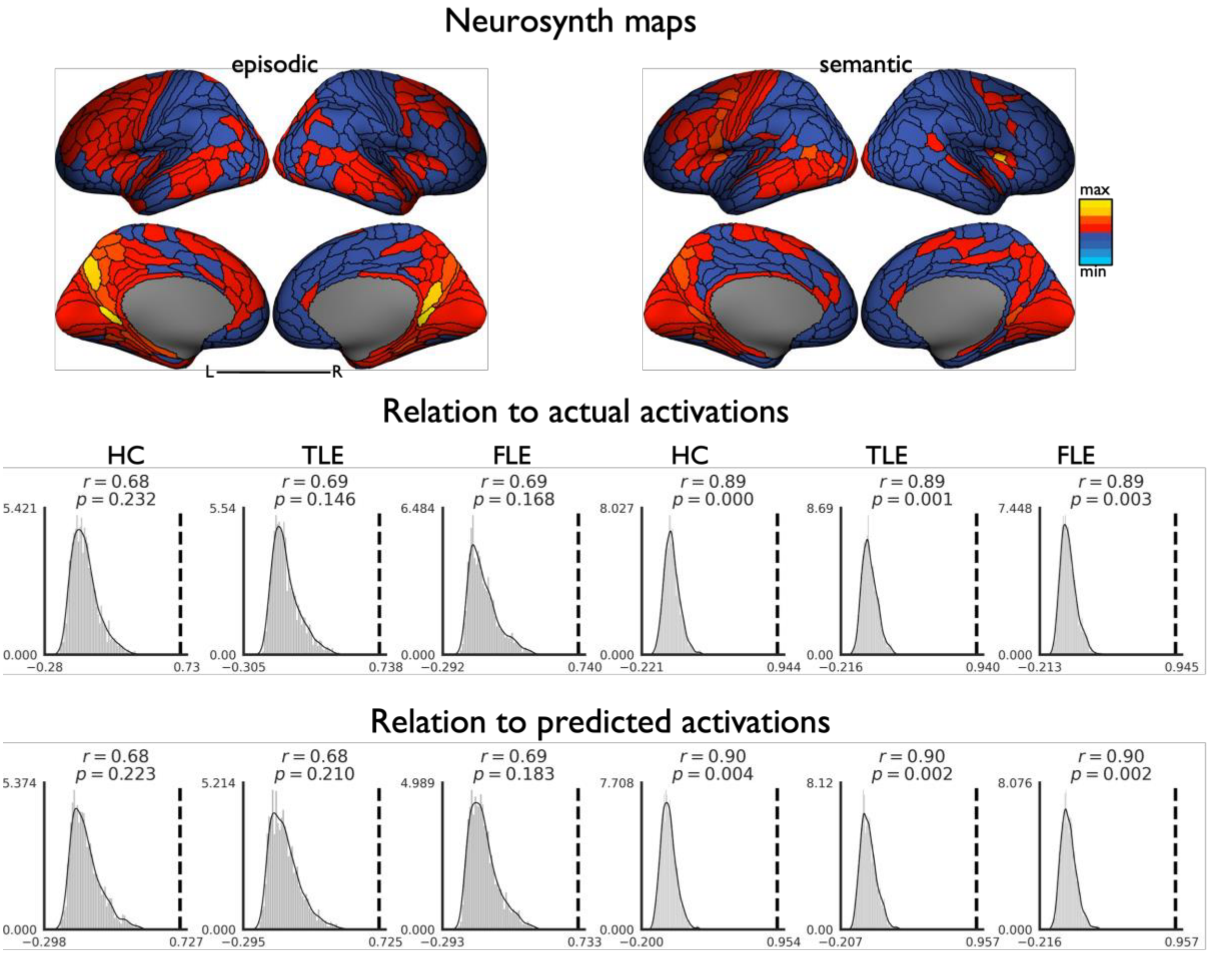


**Supplementary Figure 2. Neurosynth decoding analysis.** Neurosynth-derived task activation maps for episodic and semantic memory (*upper panel*) and their relation to actual and predicted task activation in all cohorts. **Abbreviations** | HC=healthy controls, TLE=temporal lobe epilepsy, FLE= frontal lobe epilepsy.

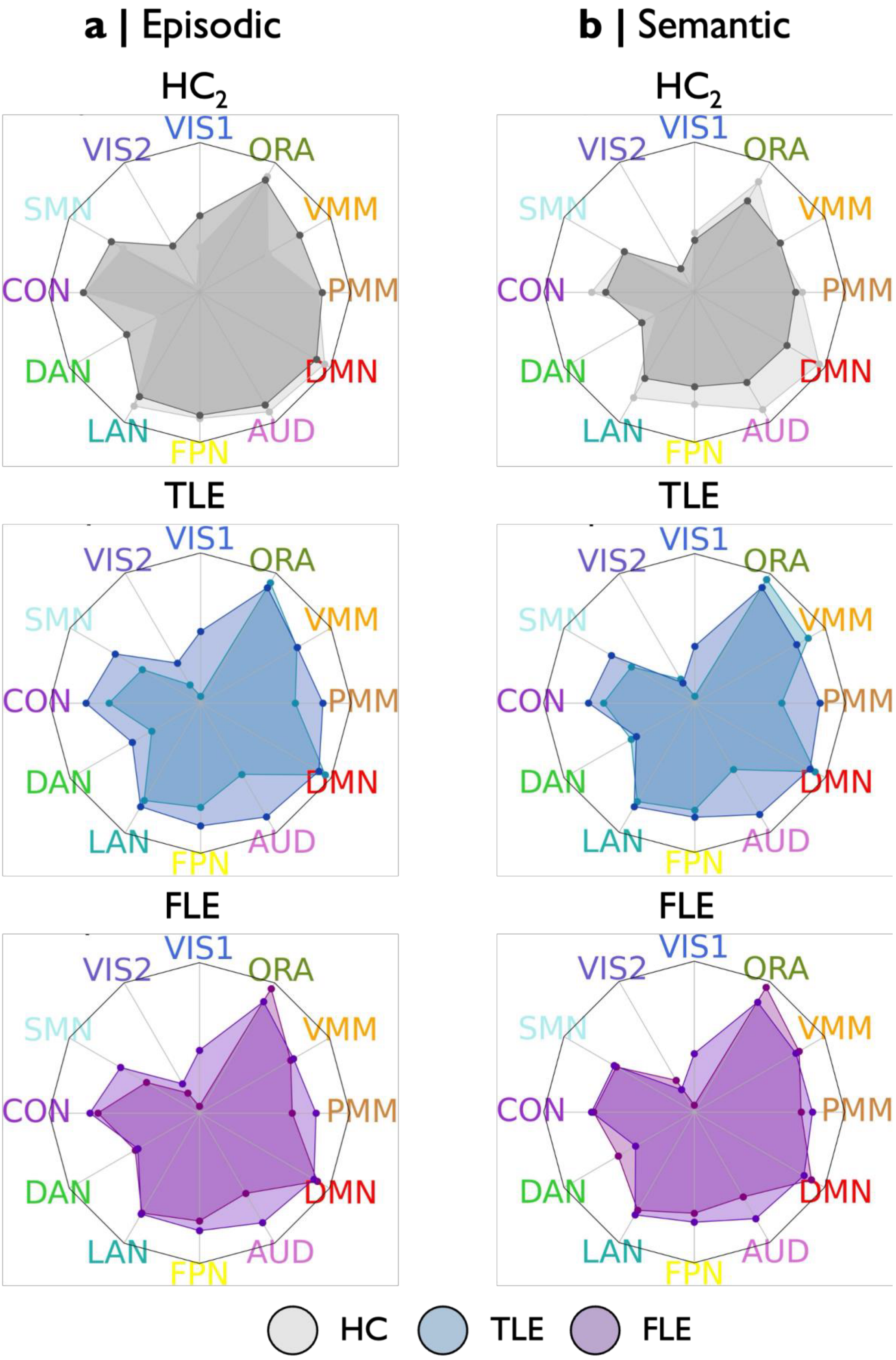
a | Episodic
b | Semantic
HC$_2$
TLE
FLE
VIS1
ORA
VMM
PMM
DMN
AUD
FPN
LAN
DAN
CON
SMN
VIS2
HC
TLE
FLE

**Supplementary Figure 3. Community-based analyses of predicted and actual task activations in each cohort.** [1,2] Darker colouring shades represent the predicted task activations, which show a broader topographical distribution compared to the actual activations (lighter shades) in both episodic **(a)** and semantic task. **Abbreviations** | HC=healthy controls, TLE=temporal lobe epilepsy, FLE= frontal lobe epilepsy, VIS1=primary visual, VIS2=secondary visual, SMN=somatomotor, CON=cingulo-opercular, DAN=dorsal attention network, LAN=language, FPN=fronto-parietal network, AUD=auditory, DMN=default mode network, PMM=posterior multimodal, VMM=ventral multimodal, ORA=orbito-affective.

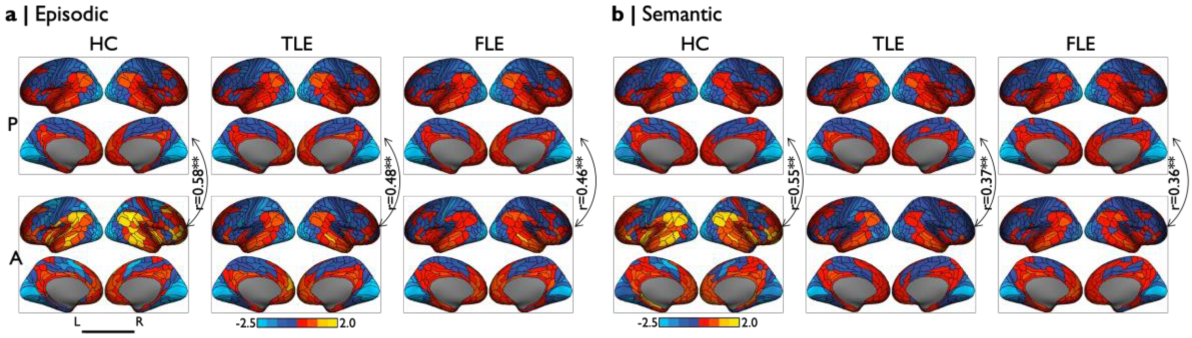


**Supplementary Figure 4. Robustness analysis using mixed task-activation template.** Predicted-to-actual activation overlap remained significant for both episodic **(a)** and semantic **(b)** tasks. Group-level activation templates used for this prediction were derived from an independent sample comprising healthy controls (n=10), TLE (n=10), and FLE (n=10) patients with variable degrees of pathology. **Abbreviations |** P=predicted activation, A= actual activation, HC=healthy controls, TLE=temporal lobe epilepsy, FLE=frontal lobe epilepsy, L=left, R=right.

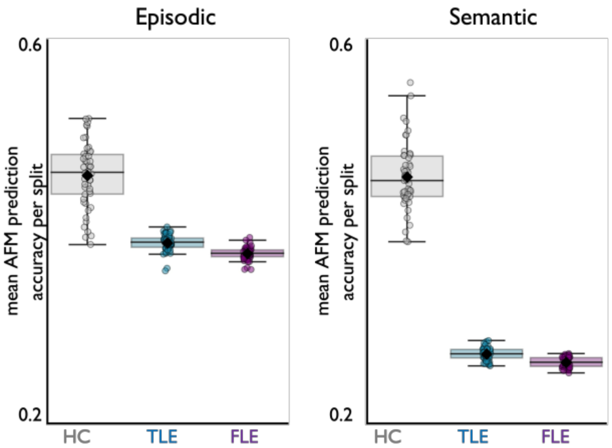


**Supplementary Figure 5. Robustness of AFM prediction accuracy across 50 random healthy-control activation template splits.** Each point represents the mean predicted-to-actual activation correlation (*r*) across participants within a group for one random split. Boxes show the distribution of prediction accuracy across the 50 random splits, and diamonds indicate the mean across splits. Results are shown separately for episodic (left) and semantic (right) retrieval. **Abbreviations** | AFM=activity flow mapping, HC=healthy controls, TLE=temporal lobe epilepsy, FLE=frontal lobe epilepsy.

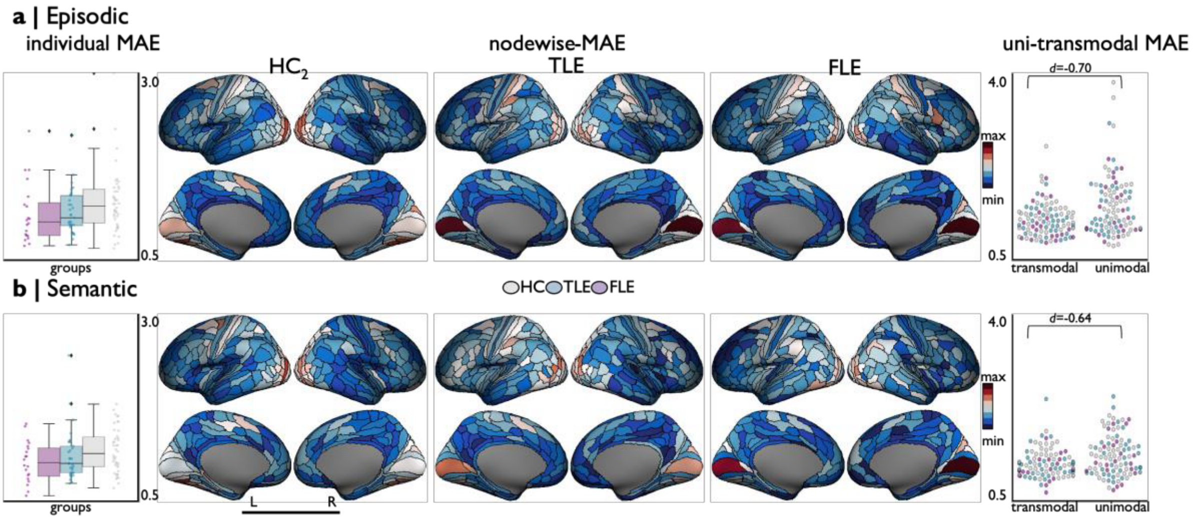


**Supplementary Figure 6**. **Mean absolute error (MAE) maps.** MAE maps derived from actual and predicted episodic **(a)** and semantic **(b)** memory task activations. Bar plots indicate individual MAE averaged for each cohort. Swarm plots indicate individual MAE for unimodal and transmodal regions. **Abbreviations |** HC=healthy controls (grey), TLE=temporal lobe epilepsy (blue), FLE=frontal lobe epilepsy (purple), L=left, R=right.

| ID | Age[+] | Sex[+] | Laterality | Diagnosis | MRI findings | HA | Resection site/Histopathology | Engel outcome |
|---|---|---|---|---|---|---|---|---|
| **TLE** | | | | | | | | |
| PX01 | 19 | F | L | mTLE | ganglioglioma | 1 | ganglioglioma, WHO grade 1 | 1a |
| PX02 | 33 | F | L | TLE | hyperintense FLAIR signal intensity in the hippocampus | 0 | hippocampal gliosis | 1a |
| PX03 | 57 | M | L | mTLE | MTS | 1 | HS | 1a |
| PX04 | 41 | F | R | mTLE | MTS | 1 | HS | 1a |
| PX05 | 28 | F | L | TLE | MTS | 1 | n/a | n/a |
| PX06 | 30 | M | L | TLE | hyperintense amygdala, which could be due to an underlying hamartoma | 0 | n/a | n/a |
| PX07 | 52 | M | L | mTLE | MTS | 1 | n/a | n/a |
| PX08 | 55 | F | L | mTLE | MTS | 0 | hippocampal FCD I | 1a |
| PX09 | 26 | F | L | mTLE | MTS | 1 | n/a | n/a |
| PX10 | 25 | M | R | mTLE | amygdala, hippocampus and temporal pole lesion with differential diagnosis of low-grade glioma vs FCD | 0 | n/a | n/a |

| | | | | | | | | |
|---|---|---|---|---|---|---|---|---|
| PX11 | 53 | F | L | mTLE | MTS | 1 | n/a | n/a |
| PX12 | 43 | M | R | mTLE | MTS | 1 | HS | 1a |
| PX13 | 32 | M | R | TLE | temporal neocortex atrophy and agenesis of piriform | 0 | n/a | 1a |
| PX14 | 27 | F | L | TLE | no clear lesion | 0 | n/a | n/a |
| PX15 | 28 | M | L | mTLE | MTS | 1 | hippocampal collagenized cavernoma | 1/a |
| PX16 | 39 | F | R | mTLE | MTS | 1 | HS | 1a |
| PX17 | 40 | F | R | TLE | no clear lesion | 1 | hippocampal gliosis | 1a |
| PX18 | 29 | F | R | mTLE | MTS | 1 | n/a | n/a |
| PX19 | 64 | M | R | TLE | no clear lesion | 0 | n/a | n/a |
| PX20 | 27 | M | L | mTLE | T2 hyperintense lesion in the left amygdala | 0 | n/a | n/a |
| PX21 | 62 | M | L | mTLE | MTS | 1 | n/a | n/a |
| PX22 | 27 | F | R | mTLE | MTS | 1 | HS | 1a |

| | | | | | | | | |
|---|---|---|---|---|---|---|---|---|
| PX23 | 42 | M | R | mTLE | amygdala lesion with differential diagnosis of low-grade glioma, glioneuronal hamartoma, or FCD | 0 | lateral neocortex subpial gliosis | 2b |
| PX24 | 22 | F | R | TLE | no clear lesion | 0 | n/a | n/a |
| PX25 | 50 | M | R | mTLE | MTS | 0 | n/a | n/a |
| PX26 | 30 | M | L | TLE | no clear lesion | 0 | n/a | n/a |
| PX27 | 32 | M | L | mTLE | MTS | 0 | n/a | n/a |
| PX28 | 30 | M | L | TLE | no clear lesion | 0 | n/a | n/a |
| PX29 | 30 | F | L | TLE | no clear lesion | 1 | n/a | n/a |
| PX30 | 44 | M | L | TLE | no clear lesion | 0 | n/a | n/a |
| PX31 | 30 | F | R | mTLE | MTS | 1 | n/a | n/a |
| PX32 | 34 | M | R | TLE | no clear lesion | 1 | n/a | n/a |
| PX33 | 28 | M | R | TLE | T2 hyperintensity in the left hippocampus | 1 | n/a | n/a |
| FLE | | | | | | | | |

| | | | | | | | | |
|---|---|---|---|---|---|---|---|---|
| PX01 | 42 | F | R | FLE-orbito-frontal | no clear lesion | 1 | frontal FCD IIA | 1a |
| PX02 | 26 | F | R | FLE-fronto-central | no clear lesion | 0 | posterior cingulate FCD IIA | 1a |
| PX03 | 27 | F | R | FLE | frontal FCD | 0 | n/a | n/a |
| PX04 | 40 | F | R | FLE- mesial frontal | T2 hyperintense lesion in the anterior right frontal region, no enhancement | 1 | frontal FCD IIA | 1b |
| PX05 | 21 | M | L | FLE | gliosis in the left fronto-parietal and temporal region | 1 | frontal FCD IIA | 1a |
| PX06 | 25 | M | R | FLE-fronto-polar | no clear lesion | 0 | n/a | n/a |
| PX07 | 19 | F | L | FLE | no clear lesion | 1 | n/a | n/a |
| PX08 | 42 | M | R | FLE | no clear lesion | 0 | n/a | n/a |
| PX09 | 18 | F | L | FLE | no clear lesion | 0 | n/a | n/a |
| PX10 | 24 | M | L | FLE | no clear lesion | 0 | n/a | n/a |
| PX11 | 54 | M | R | FLE-SSMA | no clear lesion | 0 | n/a | n/a |
| PX12 | 25 | F | L | FLE-orbito-frontal | no clear lesion | 0 | n/a | n/a |

| | | | | | | | | |
|---|---|---|---|---|---|---|---|---|
| PX13 | 22 | M | L | FLE | no clear lesion | 0 | n/a | n/a |
| PX14 | 42 | F | L | FLE | nonspecific T2/FLAIR hyperintensity in the frontal lobe | 0 | n/a | n/a |
| PX15 | 31 | F | L | FLE | no clear lesion | 0 | n/a | n/a |
| PX16 | 33 | M | R | FLE-rolandic area | increased FLAIR signal with blurring of the gray-white matter junction along right middle frontal gyrus, in keeping with FCD | 0 | n/a | n/a |
| PX17 | 23 | M | L | FLE-orbito-frontal | focal cortical developmental abnormality at the left basal frontal region and in the posterior left frontal orbital gyrus | 0 | n/a | n/a |
| PX18 | 53 | M | L | FLE-orbito-frontal | no clear lesion | 0 | n/a | n/a |
| PX19 | 29 | M | R | FLE | no clear lesion | 0 | n/a | n/a |
| PX20 | 34 | F | R | FLE-fronto-central | inferior right postcentral gyrus FCD | 0 | n/a | n/a |

**Supplementary Table 1. Demographic and clinical details.** [+]No significant age differences (one-way analysis of variance ANOVA: $F(2,137)=2.38$, $P=0.10$) and no significant differences in sex distribution across groups ($\chi2(2)=0.12$, $P=0.94$). **Abbreviations** | PX=patients, M=male, F=female, L=left lateralized, R=right lateralized, mTLE=mesiotemporal temporal lobe epilepsy, MTS=mesiotemporal sclerosis, HS=hippocampal sclerosis, FLE=frontal lobe epilepsy, FCD=focal cortical dysplasia, HA=hippocampal atrophy.

| **AFM results, Behavioral and clinical scores** | | | |
|---|---|---|---|
| **Scores** | **$HC_2$** (*n*=42) | **TLE** (*n*=33) | **FLE** (*n*=20) |
| **AFM accuracy (mean *r* ± SD, range)** | | | |
| Episodic | 0.60±0.210 | 0.44±0.26 | 0.43±0.26 |
| Semantic | 0.57±0.25 | 0.33±0.26 | 0.33±0.28 |
| **Task-specificity (*t-values*, [CI] of Fisher-*z* transformed *r*, *(effect size)*)** | | | |
| Episodic | 15.87[0.06,0.08](2.45)*** | 40.57[0.16,0.18](7.06)*** | 14.41[0.10,0.14](3.22) *** |
| Semantic | 21.01[0.07,0.08](3.24)*** | -27.92[-0.11,0.10](4.86)*** | -9.53[-0.10,-0.06](-2.13)*** |
| **Cohort-specificity (*t-values*, [CI] of Fisher-*z* transformed *r*)** | | | |
| Episodic | - | 2.78[0.004,0.03](0.48) * | −2.06[-0.3,0.0002](-0.46) |
| Semantic | - | −6.50[-0.04,-0.02](-1.13) *** | 5.10[0.02,0.05](1.14) ** |
| **Behavioral and clinical scores (mean ± SD, range)** | | | |
| Episodic Accuracy | 70.16±21.33 (12.50-100) | 45.89±21.12 (7.10-78.60) | 46.61±19.71 (19.60-94.60) |
| RTs | 1.43±0.25 (0.43-1.82) | 1.37±0.29 (0.52-1.79) | 1.35±0.40 (0.48-1.80) |
| Semantic Accuracy | 80.25±11.72 (28.60-92.90) | 77.16±13.67 (26.80-94.60) | 75.00±18.03 (23.20-91.10) |
| RTs | 1.34±0.16 (0.75-1.71) | 1.36±0.24 (0.54-1.66) | 1.32±0.26 (0.39-1.61) |
| EpiTrack | 35.83±2.73 (31-40) | 32.27±3.48 (21-40) | 31.70±5.16 (20-39) |
| Seizure Duration (years) | - | 12.64±10.60 (1-45) | 19.35±13.23 (3-46) |
| **AFM relation to MRI features, behavior and clinical scores** | | | |
| AFM accuracy and Behavioral scores (*r* [LOO]) $HC_2$/TLE/FLE (*n*=42/33/20) | | AFM accuracy and MRI features (*r* [LOO] ***(effect size)***) $HC_2$/TLE/FLE (*n*=42/33/20) | |
| Episodic | 0.18 [0.14, 0.23] | Thickness - Episodic | 0.04 [0.004, 0.13] (0.07) |
| | | - Semantic | 0.17 [0.10, 0.22] (0.34) |
| Semantic | 0.21 [0.16, 0.24] | qT1 - Episodic | 0.34 [0.31, 0.37] (0.72) ** |
| | | - Semantic | 0.16 [0.13, 0.20] (0.33) |
| EpiTrack – Episodic | 0.08 [0.05, 0.12] | SC-DC - Episodic | 0.08 [0.06, 0.14] (0.17) |
| – Semantic | 0.34 [0.27, 0.38] ** | - Semantic | 0.21 [0.14, 0.27] (0.42) |
| Seizure duration | | FC-DC - Episodic | -0.31 [-0.34, -0.21] (-0.64) * |
| – Episodic | -0.29 [-0.33, -0.23] * | - Semantic | -0.24 [-0.28, -0.17] (-0.49) * |
| – Semantic | -0.35 [-0.40, -0.29] * | FC neo-hippo Episodic | -0.21 [-0.25, -0.10] (-0.43) |
| | | - Semantic | -0.14 [-0.18, -0.05] (-0.28) |
| **Exploratory Analyses** | | | |
| | TLE (*n*=33) | TLE with HA (*n*=18) | TLE without HA (*n*=15) |
| Hippocampal volume and AFM (*r* [LOO]) | | | |
| Episodic | 0.35 [0.29, 0.42] | 0.35 [0.29, 0.45] | 0.22 [0.04, 0.33] |
| Semantic | 0.11 [0.02, 0.21] | 0.14 [0.04, 0.28] | -0.10 [-0.29, 0.07] |
| ASM and AFM (*r* [LOO]) TLE/FLE (*n*=33/20) | | | |
| Episodic | -0.24 [-0.27, -0.19] | | |
| Semantic | -0.15 [-0.20, -0.12] | | |
| **Sensitivity Analyses** | | | |
| mean tSNR and AFM $HC_2$/TLE/FLE (*n*=42/33/20) | | mean FD and AFM $HC_2$/TLE/FLE (*n*=42/33/20) | |

| | | |
|---|---|---|
| Episodic | 0.11 [0.10, 0.15] | -0.12 [-0.15, -0.09] |
| Semantic | 0.19 [0.16, 0.22] | -0.14 [-0.19, -0.11] |

**Supplementary Table 2. Activity flow mapping prediction accuracy, behavior, clinical measures, and associations with MRI-derived features.** Mean (± SD) behavioral and clinical measures are reported for healthy controls ($HC_2$), temporal lobe epilepsy (TLE), and frontal lobe epilepsy (FLE), including episodic and semantic memory accuracy, reaction times (RT), EpiTrack scores, and seizure duration. AFM prediction accuracy, task-specificity, cohort-specificity, and correlations between AFM prediction accuracy and behavioral, clinical, and MRI-derived measures are also shown. Exploratory analyses include associations with hippocampal volume and anti-seizure medication (ASM) burden, and sensitivity analyses examine relationships with resting-state temporal signal-to-noise ratio (tSNR) and framewise displacement (FD). Values are presented as mean ± SD, *t*-values with 95% confidence intervals (CI) of Fisher *z*-transformed correlation coefficients, or Pearson correlation coefficients (*r*) with leave-one-out (LOO) confidence intervals, as appropriate. Statistical significance as follows: *$P<0.05$; **$P<0.005$. ***$P<0.000001$; no asterisk indicates a non-significant result. Exact *p*-values for the main analyses are reported in the main text, and those supplementary analyses are reported in the Supplementary materials. For the correlation analyses, false discovery rate (FDR) correction was applied separately for each MRI, behavioral, and clinical measures across the episodic and semantic prediction tasks ($P_{FDR}<0.05$). The same FDR correction was applied to exploratory analyses examining associations between AFM prediction accuracy and hippocampal volume and anti-seizure medication (ASM) burden. Effect sizes are reported as Pearson's *r* for correlation analyses, and *Cohen's dz* for two-tailed paired-sample *t*-tests. **Abbreviations** | CI=confidence interval, LOO=leave-one-out confidence interval, HA=hippocampal atrophy, SC-DC=structural connectivity degree centrality, FC-DC=functional connectivity degree centrality, FC neo-hippo=neocortical–hippocampal functional connectivity, tSNR=temporal signal-to-noise ratio, FD=framewise displacement.

| Association | *r* | *p* | Bootstrap 95% CI | LOO range |
|---|---|---|---|---|
| **Episodic** | | | | |
| AFM accuracy & memory scores | -0.27 | 0.26 | [-0.65, 0.26] | [-0.34, 0.02] |
| AFM accuracy & EpiTrack | -0.22 | 0.35 | [-0.59, 0.302] | [-0.34, -0.13] |
| AFM accuracy & seizure duration | -0.14 | 0.54 | [-0.62, 0.14] | [-0.29, -0.08] |
| **Semantic** | | | | |
| AFM accuracy & memory scores | 0.21 | 0.37 | [-0.22, 0.65] | [0.03, 0.34] |
| AFM accuracy & EpiTrack | 0.41 | 0.07 | [-0.16, 0.78] | [0.21, 0.57] |
| AFM accuracy & seizure duration | -0.20 | 0.39 | [-0.60, 0.15] | [-0.31, -0.16] |

**Supplementary Table 3. Robustness of the clinical and behavioral correlations in the FLE cohort. Abbreviations** | AFM=activity flow mapping, CI=confidence interval, LOO=leave-one-out confidence interval.

**Resting-state FC changes in TLE**

| Regions A and Network Assignments | | Regions B and Network Assignments | |
|---|---|---|---|
| **Regions A** | **Net** | **Regions B** | **Net** |
| Medial Superior Temporal Area | 2 | Medial Superior Temporal Area | 2 |
| Third Visual Area | 2 | Fourth Visual Area | 2 |
| Fourth Visual Area | 2 | Third Visual Area | 2 |
| Fourth Visual Area | 2 | Area IFSa | 4 |
| Primary Auditory Cortex | 8 | VentroMedial Visual Area 2 | 2 |
| PreCuneus Visual Area | 10 | ParaHippocampal Area 3 | 5 |
| Area 7m | 9 | Area 7m | 9 |
| Area 7m | 9 | Area PF Complex | 4 |

**Resting-state FC changes in FLE**

| Regions A and Network Assignments | | Regions B and Network Assignments | |
|---|---|---|---|
| **Regions A** | **Net** | **Regions B** | **Net** |
| Sixth Visual Area | 2 | Area posterior 24 | 4 |
| Third Visual Area | 2 | Fourth Visual Area | 2 |
| Fourth Visual Area | 2 | Third Visual Area | 2 |
| Area Lateral Occipital 2 | 2 | Area 31a | 9 |
| Medial Area 7P | 7 | Area 23d | 9 |
| Area 2 | 3 | Area 43 | 4 |
| Anterior 24 prime | 4 | Area anterior 10p | 7 |
| Area a24 | 9 | Area 9 Posterior | 9 |

| | | | | | | | |
|---|---|---|---|---|---|---|---|
| Area 5L | 3 | Area posterior 10p | 7 | Area 8Ad | 9 | Area 9 anterior | 9 |
| Dorsal Area 24d | 3 | Ventral Area 6 | 3 | Area 9 Posterior | 9 | Area a24 | 9 |
| Ventral Area 24d | 3 | Area 6m anterior | 4 | Area 9 Posterior | 9 | Area anterior 32 prime | 4 |
| Ventral IntraParietal Complex | 2 | Area STSv anterior | 9 | Area 44 | 6 | Area posterior 47r | 7 |
| Area 1 | 3 | Area 3a | 3 | Area 9 anterior | 9 | Area 8Ad | 9 |
| Area 3a | 3 | Area 1 | 3 | Area 6 anterior | 5 | Area 8Av | 9 |
| Area dorsal 32 | 9 | Area 47l (47 lateral) | 9 | Entorhinal Cortex | 9 | Area 9 Posterior | 9 |
| Area 45 | 6 | Area anterior 32 prime | 4 | Area TE1 posterior | 7 | Area PFm Complex | 7 |
| Area 47l (47 lateral) | 9 | Area dorsal 32 | 9 | Area TemporoParietoOcci pital Junction 2 | 10 | Area Lateral Occipital 3 | 2 |
| Area 10v | 9 | Posterior Insular Area 2 | 4 | Area TemporoParietoOcci pital Junction 3 | 10 | RetroSplenial Complex | 7 |
| Inferior 6-8 Transitional Area | 7 | Area 25 | 9 | Area PFm Complex | 7 | Area TE1 posterior | 7 |
| Area OP4/PV | 3 | RetroInsular Cortex | 3 | Area Lateral Occipital 3 | 2 | Area TemporoParietoOcci pital Junction 2 | 10 |
| Hippocampus | 9 | Auditory 5 Complex | 6 | Area 31a | 9 | Area Lateral Occipital 2 | 2 |
| Area TG dorsal | 9 | Auditory 4 Complex | 8 | Area posterior 47r | 7 | Area 44 | 6 |
| Area PGp | 5 | RetroSplenial Complex | 7 | Area posterior 24 | 4 | Sixth Visual Area | 2 |
| Area V3CD | 2 | Area TE2 anterior | 9 | Eighth Visual Area | 2 | Medial IntraParietal Area | 5 |
| VentroMedial Visual Area 2 | 2 | Primary Auditory Cortex | 8 | RetroSplenial Complex | 7 | Area TemporoParietoOcci pital Junction 3 | 10 |
| posterior OFC Complex | 12 | posterior OFC Complex | 12 | Area 23d | 9 | Medial Area 7P | 7 |
| Area anterior 32 prime | 4 | Area 45 | 6 | Area 5m | 3 | Area PF opercular | 4 |
| Medial Superior Temporal Area | 2 | Medial Superior Temporal Area | 2 | Medial IntraParietal Area | 5 | Eighth Visual Area | 2 |
| RetroSplenial Complex | 7 | Area PGp | 5 | Area 8BM | 7 | Auditory 4 Complex | 8 |
| Seventh Visual Area | 2 | Area TE2 posterior | 5 | Area 8Av | 9 | Area 6 anterior | 5 |
| Area Lateral Occipital 1 | 2 | ParaHippocampal Area 2 | 9 | Area IFJp | 7 | PreSubiculum | 9 |
| Area 7m | 9 | Area 7m | 9 | Area 46 | 4 | Area IntraParietal 2 | 7 |
| Area 6m anterior | 4 | Ventral Area 24d | 3 | Area anterior 10p | 7 | Anterior 24 prime | 4 |
| Ventral Area 6 | 3 | Dorsal Area 24d | 3 | PreSubiculum | 9 | Area IFJp | 7 |
| Area 8Ad | 9 | Area STSv posterior | 9 | Area IntraParietal 2 | 7 | Area 46 | 4 |
| Area 47l (47 lateral) | 9 | Area Frontal Opercular 5 | 4 | Area IntraParietal 1 | 7 | Area TE1 posterior | 7 |
| Area IFSa | 4 | Fourth Visual Area | 2 | Area PF opercular | 4 | Area 5m | 3 |
| Area 9-46d | 4 | Inferior 6-8 Transitional Area | 7 | Area PF Complex | 4 | Area posterior 10p | 7 |
| Area 9-46d | 4 | Lateral Belt Complex | 8 | Area Lateral Occipital 3 | 2 | IntraParietal Sulcus Area 1 | 2 |
| Area 9 anterior | 9 | Area 10v | 9 | Area posterior 10p | 7 | Area PF Complex | 4 |
| Area 10v | 9 | Area 9 anterior | 9 | Auditory 4 Complex | 8 | Area 8BM | 7 |
| Inferior 6-8 Transitional Area | 7 | Area 9-46d | 4 | Primary Motor Cortex | 3 | Area OP4/PV | 3 |
| Area OP4/PV | 3 | Area OP1/SII | 3 | Medial Area 7A | 4 | VentroMedial Visual Area 2 | 2 |
| Area OP1/SII | 3 | Area OP4/PV | 3 | Area 2 | 3 | Supplementary and Cingulate Eye Field | 4 |
| RetroInsular Cortex | 3 | Area OP4/PV | 3 | Area 47m | 9 | Area 8BM | 7 |
| Posterior Insular Area 2 | 4 | Area 10v | 9 | Area 8B Lateral | 9 | Area STSd posterior | 6 |

| | | | | | | | |
|---|---|---|---|---|---|---|---|
| Anterior Agranular Insula Complex | 12 | Area IntraParietal 0 | 5 | Area 9 Posterior | 9 | Area PF Complex | 4 |
| Auditory 5 Complex | 6 | Hippocampus | 9 | Area IFJa | 6 | Area TE1 Middle | 9 |
| ParaHippocampal Area 3 | 5 | PreCuneus Visual Area | 10 | RetroInsular Cortex | 8 | Area PF opercular | 4 |
| Area STSv posterior | 9 | Area 8Ad | 9 | Anterior IntraParietal Area | 5 | PreSubiculum | 9 |
| Area TE2 anterior | 9 | Area V3CD | 2 | Entorhinal Cortex | 9 | PreSubiculum | 9 |
| Area TE2 posterior | 5 | Seventh Visual Area | 2 | Entorhinal Cortex | 9 | Entorhinal Cortex | 9 |
| Area IntraParietal 0 | 5 | Anterior Agranular Insula Complex | 12 | PreSubiculum | 9 | Anterior IntraParietal Area | 5 |
| Area PF Complex | 4 | Area 7m | 9 | PreSubiculum | 9 | Entorhinal Cortex | 9 |
| Area PFm Complex | 7 | Auditory 4 Complex | 8 | PreSubiculum | 9 | PreSubiculum | 9 |
| VentroMedial Visual Area 1 | 2 | Area Lateral Occipital 3 | 2 | Area STGa | 6 | Area IFJp | 7 |
| VentroMedial Visual Area 3 | 2 | Area 31a | 7 | Area STSd anterior | 6 | Area TemporoParietoOcci pital Junction 1 | 6 |
| ParaHippocampal Area 2 | 9 | Area Lateral Occipital 1 | 2 | Area TemporoParietoOcci pital Junction 1 | 6 | Area STSd anterior | 6 |
| Area Lateral Occipital 3 | 2 | VentroMedial Visual Area 1 | 2 | Area IntraParietal 2 | 7 | Area STSv anterior | 9 |
| VentroMedial Visual Area 2 | 2 | Auditory 4 Complex | 8 | Area PF opercular | 4 | RetroInsular Cortex | 8 |
| Area 31a | 7 | VentroMedial Visual Area 3 | 2 | VentroMedial Visual Area 2 | 2 | Medial Area 7A | 4 |
| Area 25 | 9 | Inferior 6-8 Transitional Area | 7 | VentroMedial Visual Area 2 | 2 | Anterior 24 prime | 4 |
| posterior OFC Complex | 12 | posterior OFC Complex | 12 | Area STSv anterior | 9 | Area IntraParietal 2 | 7 |
| Insular Granular Complex | 3 | ProStriate Area | 1 | Area TE1 Middle | 9 | Area IFJa | 6 |
| Area Frontal Opercular 5 | 4 | Area 47l (47 lateral) | 9 | Para-Insular Area | 4 | Area 8Av | 9 |
| Area posterior 10p | 7 | Area 5L | 3 | Fourth Visual Area | 2 | Area 13l | 7 |
| Lateral Belt Complex | 8 | Area 9-46d | 4 | Middle Temporal Area | 2 | Area anterior 9-46v | 7 |
| Auditory 4 Complex | 8 | Area TG dorsal | 9 | Superior Temporal Visual Area | 10 | Area posterior 10p | 7 |
| Auditory 4 Complex | 8 | VentroMedial Visual Area 2 | 2 | Area 8BM | 7 | ProStriate Area | 1 |
| Medial Superior Temporal Area | 2 | Dorsal Transitional Visual Area | 1 | Area 8Av | 9 | Para-Insular Area | 4 |
| Fourth Visual Area | 2 | Posterior Insular Area 2 | 4 | Area 9 Middle | 9 | Inferior 6-8 Transitional Area | 7 |
| Primary Motor Cortex | 3 | Area OP4/PV | 3 | Area IFSp | 7 | Area IFSa | 4 |
| Parieto-Occipital Sulcus Area 2 | 7 | Area V4t | 2 | Area IFSa | 4 | Area IFSp | 7 |
| Posterior InferoTemporal complex | 2 | Area V3CD | 2 | Area anterior 9-46v | 7 | Middle Temporal Area | 2 |
| Superior Temporal Visual Area | 6 | Area 6m anterior | 4 | Area 13l | 7 | Fourth Visual Area | 2 |
| Area 2 | 3 | Area 6mp | 3 | Inferior 6-8 Transitional Area | 7 | Area 9 Middle | 9 |
| Anterior 24 prime | 4 | Area PF opercular | 4 | Area OP4/PV | 3 | Primary Motor Cortex | 3 |
| Area IFJp | 7 | Area STGa | 6 | Anterior IntraParietal Area | 5 | PreSubiculum | 9 |
| Area IFSa | 7 | Area IntraParietal 1 | 7 | Entorhinal Cortex | 9 | Entorhinal Cortex | 9 |
| RetroInsular Cortex | 8 | Area PF opercular | 4 | Entorhinal Cortex | 9 | PreSubiculum | 9 |
| Anterior Ventral Insular Area | 7 | Area TE2 posterior | 5 | PreSubiculum | 9 | PreSubiculum | 9 |
| Frontal OPercular Area 1 | 4 | Area TA2 | 8 | PreSubiculum | 9 | Anterior IntraParietal Area | 5 |
| Anterior IntraParietal Area | 5 | PreSubiculum | 9 | PreSubiculum | 9 | Entorhinal Cortex | 9 |

| | | | | | | | |
|---|---|---|---|---|---|---|---|
| Entorhinal Cortex | 9 | PreSubiculum | 9 | ProStriate Area | 1 | Area 8BM | 7 |
| PreSubiculum | 9 | Anterior IntraParietal Area | 5 | Area STSd posterior | 6 | Area 8B Lateral | 9 |
| PreSubiculum | 9 | Entorhinal Cortex | 9 | Area PF Complex | 4 | Area 9 Posterior | 9 |
| PreSubiculum | 9 | Area STSv posterior | 9 | Area posterior 10p | 7 | Superior Temporal Visual Area | 10 |
| PreSubiculum | 9 | PreSubiculum | 9 | | | | |
| Area STGa | 6 | Area IFJp | 7 | | | | |
| Area STSv posterior | 9 | PreSubiculum | 9 | | | | |
| Area TG dorsal | 9 | Area TE1 posterior | 7 | | | | |
| Area TE1 posterior | 7 | Area TG dorsal | 9 | | | | |
| Area TE2 posterior | 5 | Anterior Ventral Insular Area | 7 | | | | |
| Area PF opercular | 4 | Anterior 24 prime | 4 | | | | |
| Area PF opercular | 4 | RetroInsular Cortex | 8 | | | | |
| Area V3CD | 2 | Posterior InferoTemporal complex | 2 | | | | |
| Area 23d | 9 | Area PGi | 9 | | | | |
| Area ventral 23 a+b | 9 | Area 10v | 9 | | | | |
| Area ventral 23 a+b | 9 | VentroMedial Visual Area 2 | 2 | | | | |
| Area 6m anterior | 4 | Superior Temporal Visual Area | 6 | | | | |
| Area posterior 9-46v | 7 | Area PF opercular | 4 | | | | |
| Area 11l | 7 | PreSubiculum | 9 | | | | |
| Area OP4/PV | 3 | Primary Motor Cortex | 3 | | | | |
| Area 52 | 8 | Medial Belt Complex | 8 | | | | |
| Posterior Insular Area 2 | 4 | Fourth Visual Area | 2 | | | | |
| Frontal OPercular Area 2 | 3 | Frontal OPercular Area 1 | 4 | | | | |
| Anterior IntraParietal Area | 5 | PreSubiculum | 9 | | | | |
| Entorhinal Cortex | 9 | PreSubiculum | 9 | | | | |
| PreSubiculum | 9 | PreSubiculum | 9 | | | | |
| PreSubiculum | 9 | Area 11l | 7 | | | | |
| PreSubiculum | 9 | Anterior IntraParietal Area | 5 | | | | |
| PreSubiculum | 9 | Entorhinal Cortex | 9 | | | | |
| Dorsal Transitional Visual Area | 1 | Medial Superior Temporal Area | 2 | | | | |
| Area IntraParietal 1 | 7 | Area 6m anterior | 4 | | | | |
| Area IntraParietal 1 | 7 | Area IFSa | 7 | | | | |
| Area PF opercular | 4 | Area posterior 9-46v | 7 | | | | |
| Area PGi | 9 | Area 23d | 9 | | | | |
| VentroMedial Visual Area 2 | 2 | Area ventral 23 a+b | 9 | | | | |
| Medial Belt Complex | 8 | Area 52 | 8 | | | | |

**Supplementary Table 4. Resting-state functional connectivity differences between healthy controls and TLE (*left panel*) and FLE (*right*) patients.** Network assignments based on Glasser parcellation[1,2] are also indicated. **Abbreviations |** FC=functional connectivity, HC=healthy controls, TLE=temporal lobe epilepsy, FLE=frontal lobe epilepsy, Network assignments: VIS1(1)=primary visual ,VIS2(2)=secondary visual, SMN(3)=somatomotor, CON(4)=cingulo-opercular, DAN(5)=dorsal attention network, LAN(6)=language, FPN(7)=fronto-parietal network, AUD(8)=auditory, DMN(9)=default mode network, PMM(10)=posterior multimodal, VMM(11)=ventral multimodal, ORA(12)=orbito-affective.

| **Analysis** | **Relation to episodic memory** *(r [effect size]* | **Relation to semantic memory** *(r [effect size]* | **Relation to EpiTrack** *(r [effect size]* | **Relation to seizure Duration** *(r [effect size]* |
|---|---|---|---|---|
| **AFM analysis** | | | | |
| Episodic AFM accuracy | | | | |
| $HC_2$/TLE/FLE | 0.18 [0.35] | -- | 0.08 [0.17] | -- |
| $HC_2$/TLE | 0.24 [0.49] | -- | 0.15 [.0.31] | -- |
| $HC_2$/FLE | 0.15 [0.30] | -- | 0.10 [0.21] | -- |
| TLE/FLE | -- | -- | -- | -0.29 [-0.60]* |
| Semantic AFM accuracy | | | | |
| $HC_2$/TLE/FLE | -- | 0.21 [0.44] | 0.34 [0.73]* | -- |
| $HC_2$/TLE | -- | 0.19 [0.40] | 0.26 [0.53] | -- |
| $HC_2$/FLE | -- | 0.31 [0.65] | 0.40 [0.86] | -- |
| TLE/FLE | -- | -- | -- | -0.35 [-0.74]* |
| **Traditional analysis** | | | | |
| Episodic task-differences | | | | |
| $HC_2$ *vs* TLE | 0.09 [0.17] | -- | 0.18 [0.36] | -- |
| $HC_2$ *vs* FLE | 0.12 [0.24] | -- | 0.04 [0.09] | -- |
| TLE/FLE | -- | -- | | -0.13 [-0.26] |
| Semantic task-differences | | | | |
| $HC_2$ *vs* TLE | -- | -0.06 [-0.11] | 0.29 [0.62]* | -- |
| $HC_2$ *vs* FLE | -- | -0.01 [0.02] | 0.17 [0.36] | -- |
| TLE/FLE | -- | -- | | 0.06 [0.12] |
| Rest-differences | | | | |
| $HC_2$ *vs* TLE | 0.15 [0.30] | 0.14 [-0.28] | 0.03 [0.06] | -- |
| $HC_2$ *vs* FLE | 0.25 [0.51] | 0.22 [0.46] | 0.12 [0.24] | -- |
| TLE/FLE | -- | -- | -- | -0.13 [-0.26] |
| FC-DC | | | | |
| HC/TLE | -0.03 [-0.07] | 0.08 [0.16] | -0.10 [-0.20] | -- |
| HC/FLE | 0.01 [0.02] | 0.14 [0.28] | -0.13 [-0.27] | -- |
| TLE/FLE | -- | -- | -- | -0.01 [-0.02] |

**Supplementary Table 5. Comparison of clinical associations between activity-flow mapping prediction accuracy and conventional task-activation and resting-state measures.** Associations with episodic and semantic memory performance, EpiTrack scores, and disease duration are reported for AFM prediction accuracy and conventional task-activation differences, resting-state functional connectivity (rsFC) abnormalities, and FC-DC. Comparisons are shown for healthy controls versus patients with TLE and FLE, as well as between the two patient groups where applicable. Statistical significance as follows: *$P$<0.05; no asterisk indicates a non-significant result. Exact *p*-values for the main analyses are reported in the main text, and those supplementary analyses are reported in the Supplementary materials. For the correlation analyses, false discovery rate (FDR) correction was applied separately for each of the traditional analyses, AFM, behavioral and clinical measures across the episodic and semantic prediction tasks ($P_{FDR}$<0.05).) Effect sizes are reported as Pearson's *r* for correlation analyses and *Cohen's d*. **Abbreviations** | AFM=activity-flow mapping, FC-DC=functional connectivity degree centrality, HC= healthy controls, TLE=temporal lobe epilepsy, FLE=frontal lobe epilepsy.
